# Nanoluciferase reporter preserves immunocompetent glioma model fidelity while facilitating longitudinal molecular imaging

**DOI:** 10.64898/2026.08.24.746894

**Authors:** Carla Bianca Luena Victorio, Wisna Novera, Arun Ganasarajah, Joanne Ong, Shantanu Gupta, Eng Eong Ooi, Sven Hans Petersen, Rasha Msallam, Ann-Marie Chacko

## Abstract

Glioblastoma studies employ syngeneic orthotopic models to preserve tumor–immune interactions, but intracranial tumor burden is challenging to monitor longitudinally. Bioluminescence imaging enables non-invasive assessment, although reporter immunogenicity may compromise model fidelity. We engineered murine GL261 glioma cells to stably express nanoluciferase (NLuc) and compared them with parental GL261 (WT) and GL261 cells expressing red-shifted firefly luciferase (Red-FLuc). *In vitro*, GL261-NLuc retained growth kinetics and morphology comparable to GL261-WT and produced >100-fold stronger bioluminescence than GL261-Red-FLuc. In immunocompetent mice, GL261-NLuc formed lethal brain tumors with survival and tumor histopathology, immune profile, and response patterns to experimental oncolytic virus therapy broadly resembling GL261-WT. In contrast, GL261-Red-FLuc tumors regressed and exhibited heightened inflammation and increased infiltration of activated CD8⁺ T-cells. Longitudinal imaging of GL261-NLuc tumors detected treatment-associated changes in growth kinetics not captured by survival alone. These establish GL261-NLuc as a practical reporter for longitudinal immunocompetent glioblastoma studies amenable to immunotherapy evaluations.

**Teaser:** GL261-NLuc bioluminescent tumor cell enables non-invasive monitoring of brain cancer treatment in mice without distorting the immune response

## Introduction

Glioblastoma multiforme (GBM) remains the most aggressive and lethal primary brain malignancy and is marked by diffuse invasion, extensive heterogeneity, and a profoundly immunosuppressive microenvironment which, collectively, limit therapeutic progress (*1, 2*). Thus, preclinical development and clinical translation rely heavily on models that recapitulate both tumor biology and host immunity (reviewed recently in (*3–5*)), particularly for immunotherapy, vaccine, and virotherapy studies. Among immunocompetent murine glioma models (*6, 7*), the syngeneic GL261 model in C57BL/6 mice is one of the most widely used, as it supports tumor growth within an intact immune system while reproducing key histopathologic and therapeutic features of high-grade glioma (*8–10*). However, its orthotopic intracranial location poses a significant limitation: tumor burden cannot be measured directly, restricting longitudinal assessment and often necessitating reliance on survival-based endpoints.

Non-invasive imaging partially addresses this challenge. While MRI (Magnetic Resonance Imaging) provides high anatomical resolution (*11, 12*), and PET/SPECT (Positron Emission Tomography/ Single-Photon Emission Computed Tomography) offers functional and molecular insight (*13–15*), these modalities are costly, low-throughput, and technically demanding. Optical imaging, particularly bioluminescence imaging (BLI), is rapid, cost-effective, and well-suited for longitudinal studies, facilitating sensitive detection of intracranial tumor growth with low background noise (*16–18*). A critical limitation of reporter-based imaging, however, is that reporter proteins can alter tumor–host interactions. Luciferase-based reporters, including firefly luciferase (Fluc/Luc2) and red-shifted variants (Red-FLuc), have been shown to induce immune activation and anti-tumor responses, compromising model fidelity (*19–23*). This is an under-appreciated problem that can confound results on therapeutic efficacy studies. Nanoluciferase (NLuc) represents an attractive alternative due to its small size (∼19 kDa vs. ∼60 kDa for Red-FLuc) and high intrinsic brightness (*24–26*), features that may reduce the biological penalty of stable reporter expression while preserving imaging performance. Although NLuc has historically been less common in brain tumor models, newer substrate systems have improved its utility for *in vivo* imaging, raising the possibility that NLuc could provide a more faithful reporter platform for orthotopic glioma studies (*27–29*).

Here, we engineered GL261 cells to stably express NLuc (GL261-NLuc) and evaluated their suitability for sensitive longitudinal imaging of brain tumors in immunocompetent C57BL/6 mice. We investigated whether GL261-NLuc preserves key molecular, histologic, and immunologic features of the parental line and compared its BLI performance for non-invasive tumor growth monitoring against GL261-Red-FLuc. Further, we tested its utility in detecting changes in tumor growth patterns following therapeutic intervention with an experimental Zika virus live-attenuated vaccine (ZIKV-LAV) repurposed as oncolytic therapy (*30, 31*). By validating GL261-NLuc in these conditions, we provide functional evidence of GL261-NLuc as a practical platform for mechanistic and therapeutic studies in GBM, which minimally relies on traditional terminal measures and reduces the number of animals required in longitudinal study designs.

## Results

### Stable NLuc preserves GL261 growth and boosts BLI sensitivity

Incucyte live cell imaging revealed all three cell lines displayed a fibroblast-like morphology (**Fig. 1A**) and exhibited growth curves with comparable doubling times (*T*_d_)—12.7 h *vs.* 12.9 h *vs.* 13.6 h in WT *vs.* NLuc *vs.* Red-FLuc, respectively—at high density seeding (9,645 cells/cm^2^) (**Fig. 1B**). T*_d_* among the three cell lines remained comparable at higher seeding densities (15,625 cells/cm^2^ or 31,250 cells/cm^2^) but not at lower seeding density (3,215 cells/cm^2^), where the GL261-Red-Fluc cells divided more slowly than the other cell lines (**Fig. S1**). Further, cell seeding at very low density (1,052 cells/cm²) required for colony formation assays resulted in GL261-NLuc cells forming 20% less cell growth than GL261-WT (*p* < 0.001) and 40% less cell growth than GL261-Red-FLuc (*p* < 0.001; **Fig. 1C**), indicating a reduction in GL261-NLuc clonogenic potential under sparse culture conditions.

**Fig. 1.**
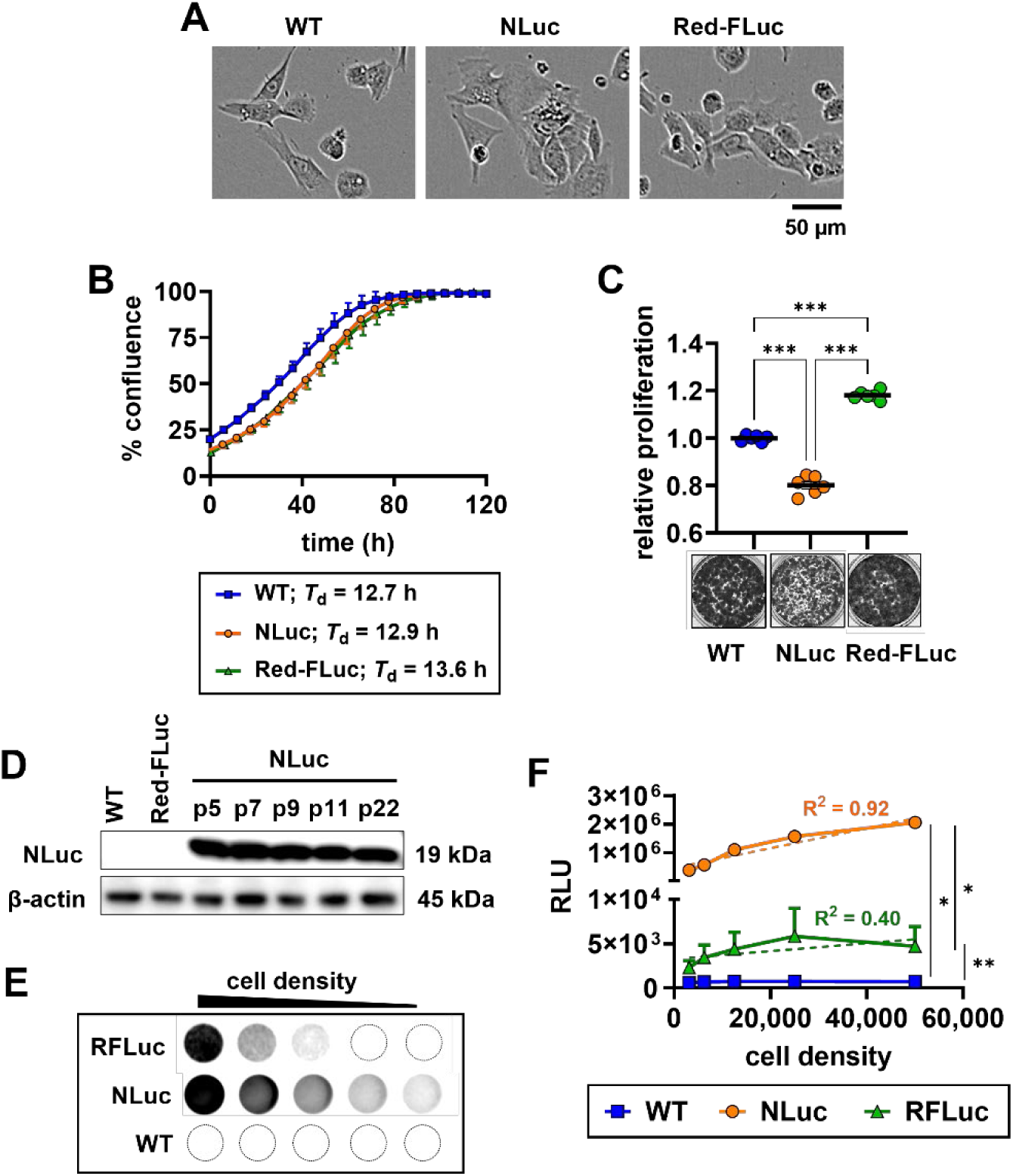
Characterization of GL261 glioma cells stably expressing NLuc. (**A**) Representative live-cell images of parental (*GL261-WT*) cells and GL261 cells expressing either red-shifted firefly luciferase (*Red-FLuc*) or nanoluciferase (*NLuc*). (**B**) Growth kinetics of cells seeded at a density of 9,645 cells/cm^2^. (**C**) Clonogenicity assay with stained colony images shown in the inset. (**D**) Western blot images of NLuc expression across passages 5, 7, 9, 11, and 22. (**E**) Representative *in vitro* bioluminescence imaging of GL261-NLuc and GL261-Red-FLuc cells at various seeding densities. Red-FLuc and NLuc signals were detected using D-luciferin and fluorofurimazine, respectively. (**F**) Quantification of luminescence. Dashed lines indicate linear regression fits; corresponding R^2^ values are shown in matching colors. **B** and **F** are shown as means ± SEM from three independent experiments. **C** is shown as mean ± SD from a representative experiment. Means in **C** and AUC in **F** were compared by Welch’s ANOVA with Dunnett’s correction. \**p* < 0.05; \*\**p* < 0.005; \*\*\**p* < 0.001; *ns*, not significant. *T_d_*, doubling time. *RLU*, relative light unit. *R^2^*, linearity coefficient.

NLuc expression in virally transduced cells was found to be stable through Western blot analysis across early and late passages (**Fig. 1D**) and even beyond passage 40 (*data not shown*); and *in vitro* bioluminescence imaging (BLI; **Fig. 1E**). NLuc signal intensity scaled linearly with cell number (R^2^ = 0.92), in contrast to Red-FLuc (R^2^ = 0.40; **Fig. 1F**). Moreover, GL261-NLuc cells produced luminescence signals > 100-fold brighter than GL261-Red-FLuc (*p* = 0.002) at matched cell densities (**Fig. 1F**).

### NLuc preserves lethality and immunosuppressive milieu of orthotopic GL261 tumors

Parental GL261-WT, GL261-NLuc, or GL261-Red-FLuc cells were orthotopically implanted into the right cerebral hemisphere (**Fig. 2A**). Mice bearing GL261-NLuc tumors developed lethal disease with median survival (OS_m_) comparable to GL261-WT (24.5 *vs* 23 days, respectively; *p* = ns) and presented with corresponding body weight loss over time due to disease (**Fig. 2B-C**). By comparison, GL261-Red-FLuc implantation did not result in lethality (OS_m_ > 30 days *vs.* GL261-WT; *p* = 0.001) or overt weight loss in progressive disease (*p* < 0.001; **Fig. 2B-C**).

**Fig. 2.**
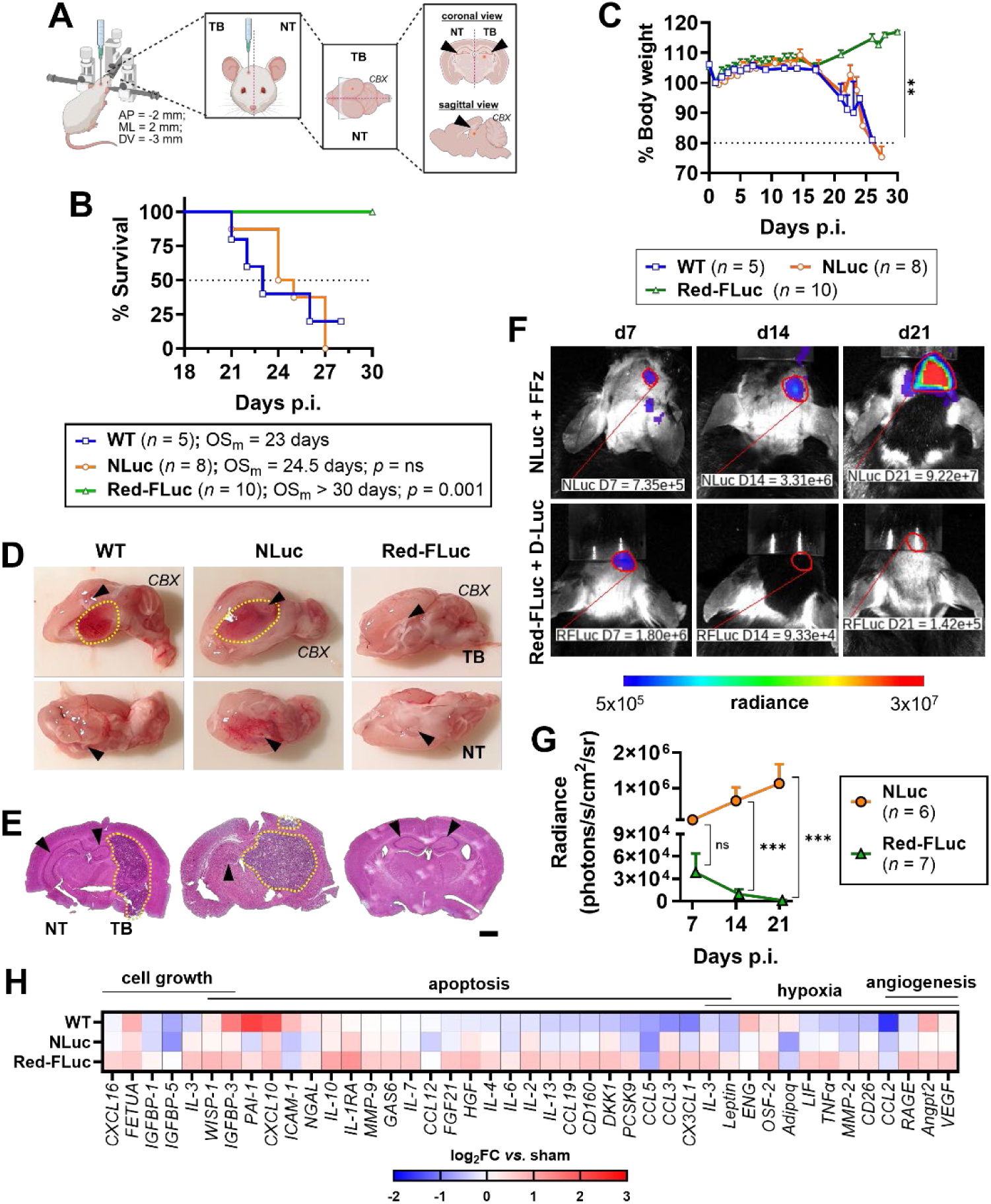
NLuc enables serial imaging of brain gliomas without altering disease progression. (**A**) Diagram of intracranial implantation of either parental GL261 (*WT*), GL261-NLuc (*NLuc)*, or GL261-Red-FLuc (*Red-FLuc*) cells into the tumor-bearing (*TB*) brain hemisphere. Injection site is depicted as red dots. (**B**) Kaplan–Meier survival curves. (**C**) Body weight change across time. (**D**-**E**) Gross pathology in sagittal view (**D**); and histopathology in coronal view (**E**) of brains. (**F**) Representative serial BLI of NLuc and Red-FLuc tumors imaged with fluorofurimazine (*FFz*) or D-luciferin (*D-Luc*), respectively. (**G**) Signal quantification of **F**. (**H**) Cytokine/chemokine heatmap in TB brains, shown as log_2_ fold change (log_2_FC) relative to sham. Median survivals (OS_m_) in **B** were compared with Mantel– Cox test with Holm–Šídák correction. AUCs in **C** were compared by one-way ANOVA with Tukey’s correction. Radiance in **G** was compared by Welch’s *t* test. \**p* < 0.05; \*\**p* < 0.005; \*\*\**p* < 0.001; *ns*, not significant. Arrows in **A**, **D**, and **E** indicate the hippocampus. Colored lines in **D-F** indicate tumor outlines. Assays in **D**, **E**, and **H** were conducted using brains collected on day 21 post-implantation (p.i.). *NT*, non-tumor-bearing; *CBX*, cerebellar cortex; *AP*, anterior-posterior; *ML*, medial-lateral; *DV*, dorsal-ventral. Scale bar = 1 mm.

Gross pathology of brains harvested from mice that succumbed at day 21 post-implantation (p.i.) revealed visible tumors in GL261-WT and GL261-NLuc tumor-bearing (TB) brain hemispheres near the hippocampus, whereas GL261-Red-FLuc-implanted brains had no visible tumor masses (**Fig. 2D**). The contralateral hemisphere without tumor served as the non-tumor-bearing (NT) control (**Fig. 2A,2D**). Histopathological analysis confirmed well-defined tumors with smooth margins in GL261-WT and GL261-NLuc groups, but not in GL261-Red-FLuc (**Fig. 2E**).

Longitudinal *in vivo* BLI revealed progressive signal increase over three weeks in GL261-NLuc tumors, whereas GL261-Red-FLuc signals were transient and declined dramatically after day 7 (*p* < 0.001 *vs.* GL261-NLuc at both days 7 and 14; **Fig. 2F,2G**).

GL261-NLuc and GL261-WT TB brains exhibited broadly similar patterns of inflammatory cytokine expression, including the downregulation of apoptosis-associated proteins and upregulation of proteins involved in cell growth (**Fig. 2H**). Both groups dampened expression of IL6, CCL5, and CX3CL1 relative to sham, reflecting an immunosuppressive phenotype. Reduced expression of hypoxia-associated proteins (*e.g.,* TNFα and MMP2) and angiogenesis-related factors (*e.g.,* CCL2 and RAGE) were similarly observed between parental and GL261-NLuc TB brains. In contrast, GL261-Red-FLuc TB brains displayed a distinct profile characterized by broad upregulation of inflammatory markers, including angiogenesis-related and apoptosis-related proteins (**Fig. 2H**).

### GL261-NLuc recapitulates the immune microenvironment of parental GL261 tumors

TB brain hemispheres were profiled by flow cytometry (FCM) at days 7 and 21 p.i. using manual gating strategy (**Fig. S2**). On day 7 the immune composition of cell-implanted TB brains, which was predominantly B and T cells, was comparable to non-tumor (NT) brain hemispheres, (**Fig. 3A**; **Fig. S3A**). On day 21, this landscape changed, becoming predominantly comprised of myeloid-derived macrophages (MDM), natural killer (NK) cells, and T cells (**Fig. 3A**). Meanwhile, NT brain immune composition remained similar to day 7 with slightly increased abundance of B and T cells (**Fig. S3A**). These findings focused our analysis on TB brains, which exhibited increased abundance of immune cells, particularly B, NK, and T lymphoid cells at day 21 relative to day 7 (**Fig. S4**).

**Fig. 3.**
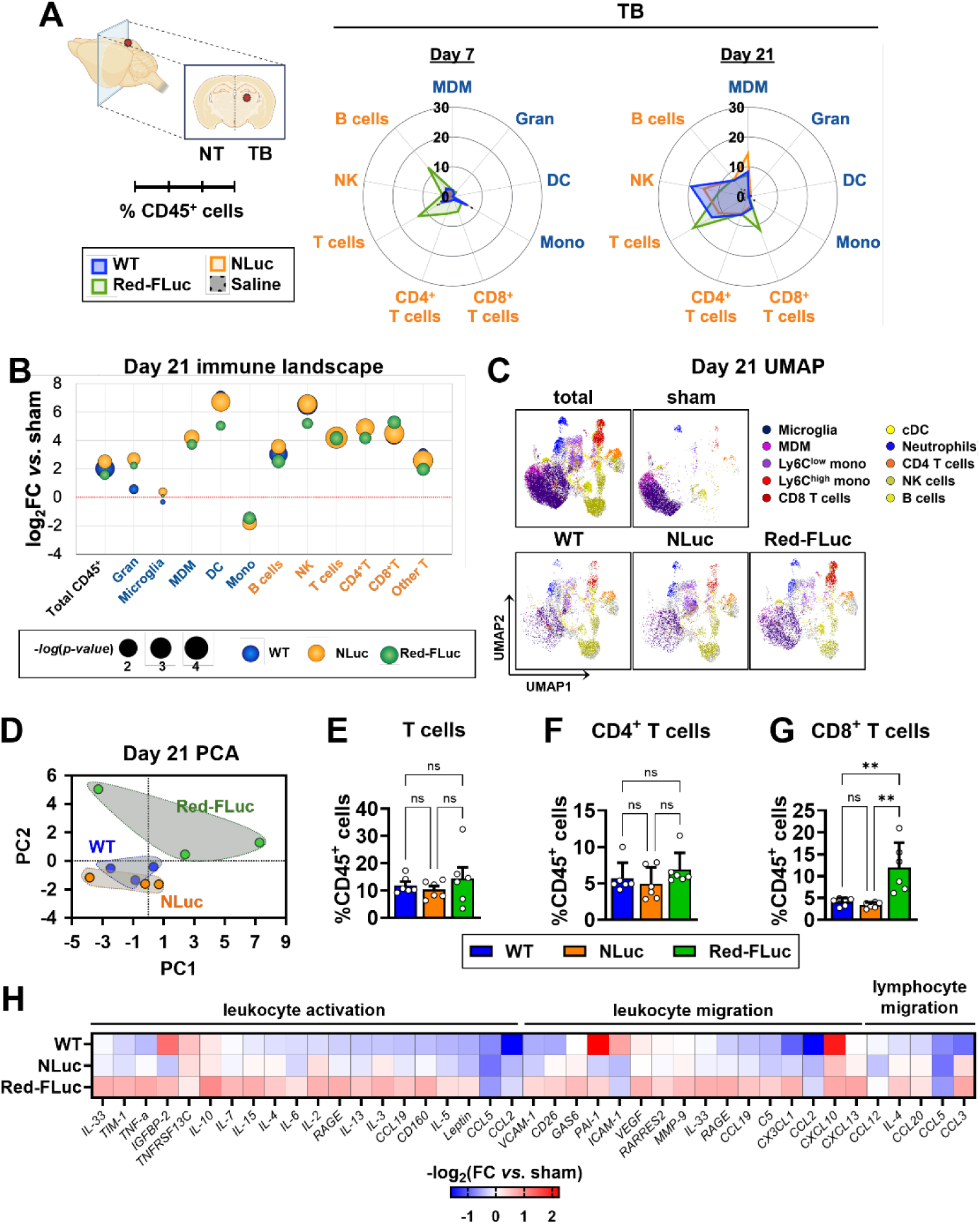
GL261-WT and GL261-NLuc gliomas exhibit comparable immune microenvironments and inflammatory profiles in male mice. (**A**) Radar plots showing immune cell populations as percentages of CD45+ cells in tumor-bearing (*TB*) brain hemispheres of mice implanted with either parental GL261 (*WT*), GL261-NLuc (*NLuc*), GL261-Red-FLuc (*Red-FLuc*), or saline (*sham*) at days 7 and 21 post-implantation (p.i.). (**B**) Bubble plot of immune composition on day 21 presented as log2 fold change (FC) relative to sham controls; bubble size is inversely proportional to *p*-value. (**C**) UMAP visualization of immune profiles in TB brains at day 21. (**D**) Principal component analysis of immune composition in TB brains at day 21. (**E-G**) Relative abundance of total T cells (**E**), CD4^+^ T cells (**F**), and CD8^+^ T cells (**G**) in TB brains shown as percentages of CD45^+^ cells. Data are mean ± SD from 6 mice per group. (**H**) Heatmap of inflammatory cytokine and chemokine expression in TB brains at day 21, shown as log_2_ FC relative to sham controls. **E-G** were analyzed by one-way ANOVA with Tukey’s correction. \**p* < 0.05; \*\**p* < 0.005; \*\*\**p* < 0.001; *ns*, not significant. *Gran*, granulocytes; *MDM*, monocyte-derived macrophages; *DC*, conventional dendritic cells; *Mono*, monocytes.

On day 7, the immune profile of GL261-NLuc TB brains closely resembled both parental and sham TB brains (**Fig. 3A**; **Fig. S5A**). By day 21, CD45^+^ immune cells in parental and GL261-NLuc TB brains were 4.0-fold (2.02 log_2_-fold) and 5.7-fold (2.51 log_2_-fold) higher than sham, respectively; and was accompanied by broad increases in myeloid and lymphoid cells except monocytes and microglia (**Fig. 3B**). Similarly increased abundance *vs.* sham in CD4^+^ T cells (26.5-fold or 4.73log_2_-fold in GL261-WT; and 29.8-fold or 4.90 log_2_-fold in GL261-NLuc) and CD8^+^ T cells (20.7-fold or 4.37 log_2_-fold in GL261-WT; and 22.7-fold or 4.50 log_2_-fold in GL261-NLuc) were noted in TB brains at day 21 (**Fig. 3B**). These findings were supported by UMAP visualization (**Fig. 3C; Fig. S5B**) and corroborated by unsupervised FlowSOM analysis (**Fig. S6**).

More importantly, principal component analysis (PCA) of all immune cell abundance clustered parental and GL261-NLuc together and separated both from GL261-Red-FLuc (**Fig. 3D**). Compared to parental and GL261-NLuc, GL261-Red-FLuc TB brains exhibited higher T cell infiltration at day 21 (**Fig. 3E-G**), most strikingly in cytotoxic CD8⁺ T cells (12.0 ± 5.7% of CD45^+^ cells) which were 2.9-fold (± 1.4; *p* = 0.003) and 3.6-fold (± 1.5; *p* = 0.001) higher than parental (4.1 ± 0.9% of CD45^+^ cells) and GL261-NLuc (3.6 ± 0.7% of CD45^+^ cells) TB brains, respectively (**Fig. 3G**). While the abundance of other lymphocytes (NK and B) in TB brains at day 21 were comparable among groups, select myeloid cells were modulated by tumor cell type implanted (**Fig. S7**). Compared to parental and GL261-NLuc TB brains, microglia abundance in GL261-Red-FLuc TB brains (29.0 ± 6.8% of CD45^+^ cells) was 160-205% higher (*p* = 0.03 for both), while conventional dendritic cell abundance (0.31 ± 0.23% of CD45^+^ cells) was 57% lower than in GL261-NLuc TB brains (0.72 ± 0.24% of CD45^+^ cells; *p* = 0.03; **Fig. S7**).

Consistent with these cellular differences, cytokine profiling relative to sham controls revealed reduced inflammatory signaling in most proteins associated with activation and migration of leukocytes in parental and GL261-NLuc TB brains (**Fig. 3H**). In contrast, GL261-Red-FLuc tumors displayed broad upregulation of these proteins (**Fig. 3H**), indicating a pro-inflammatory state, whereas GL261-NLuc retains the immunosuppressive tumor milieu of the parental model.

### NLuc preserves the T cell landscape of parental GL261 tumors

T cell status in TB and NT brains were also profiled by FCM on days 7 and 21 p.i. Across all groups, a shift from naïve (CD62L⁺CD44⁻) to effector memory (EM; CD62L⁻CD44⁺) and central memory (CM; CD62L⁺CD44⁺) state among CD4^+^ and CD8^+^ T cell populations was observed in TB brains from day 7 to day 21 (**Fig. 4A**; **Fig. S8**). On the other hand, contralateral NT brains remained enriched in naïve T cells (**Fig. S3B**).

**Fig. 4.**
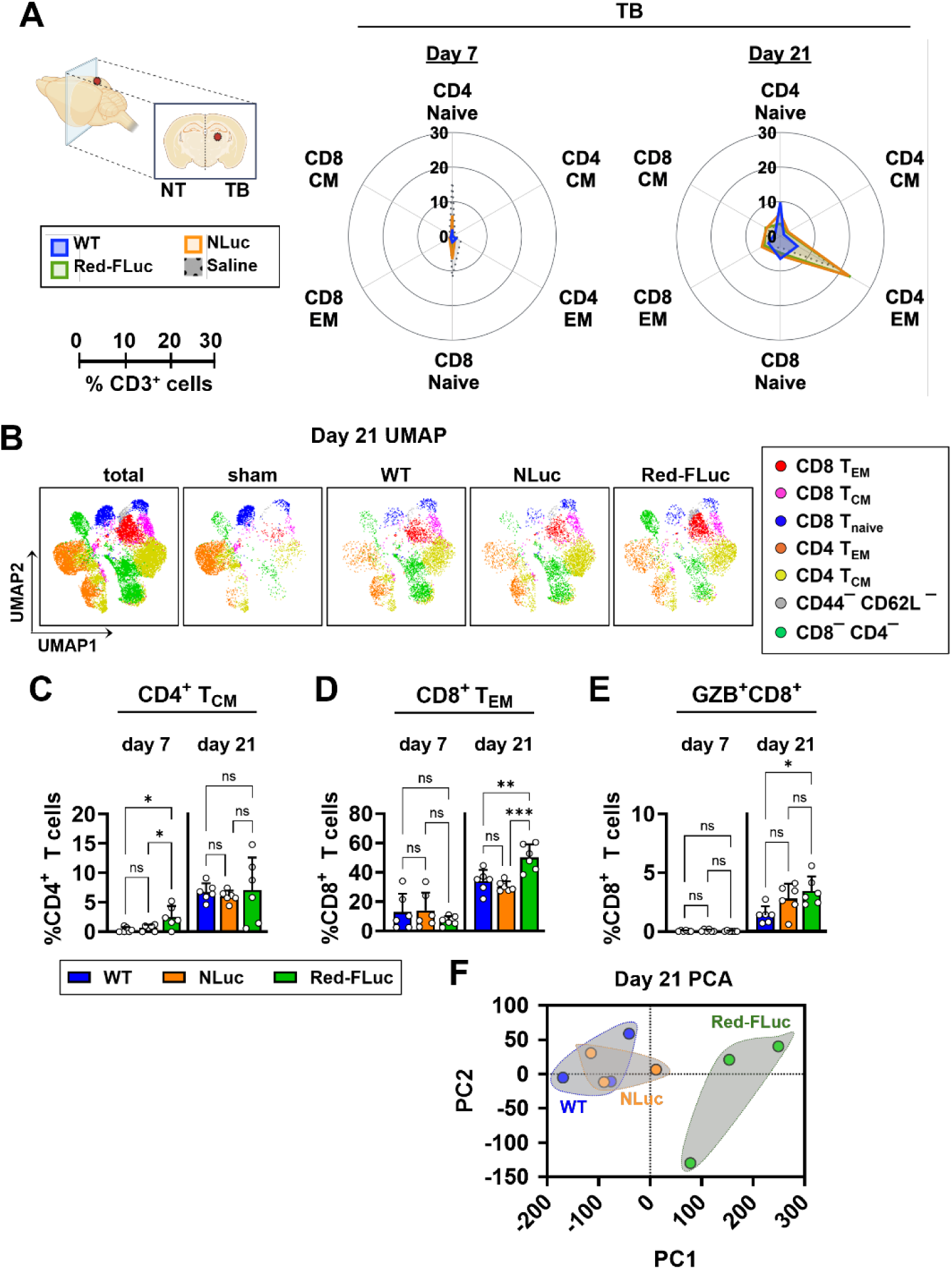
GL261-WT and GL261-NLuc gliomas exhibit comparable T cell landscapes in male mice. (**A**) Radar plots showing T cell populations as percentages of CD3+ cells in tumor-bearing (*TB*) brain hemispheres of mice implanted with either parental GL261 (*WT*), GL261-NLuc (*NLuc*), GL261-Red-FLuc (*Red-FLuc*), or saline (*sham*) at days 7 and 21 post-implantation. (**B**) UMAP visualization of the T cell landscape at day 21. (**C**-**E**) Relative abundance of CD4^+^ central memory T (T_CM_) cells (**C**), CD8^+^ effector memory T (T_EM_) cells (**D**), and granzyme B–positive (GZB^+^) CD8^+^ T cells at days 7 and 21 (**E**). (**F**) Principal component analysis (PCA) of T cell composition in TB brains on day 21. Data in **C-E** are shown as mean ± SD from 6 mice per group and analyzed by one-way ANOVA with Tukey’s correction. \**p* < 0.05; \*\**p* < 0.005; \*\*\**p* < 0.001; *ns*, not significant. *NT,* non tumor-bearing.

Whereas UMAPs at day 7 revealed highly similar T cell profiles among tumor-implanted and sham TB brains (**Fig. S9**), day 21 UMAPs highlighted the increase in CD8^+^ T_EM_ and CD4^+^ T_EM_ relative to sham particularly for GL261-Red-FLuc TB brains (**Fig. 4B**). The abundance of T_naive_, T_CM_, and T_EM_ cells among CD4^+^ and CD8^+^ T cells in TB brains was comparable between parental and GL261-NLuc groups at both timepoints, but those of CD4^+^ T_CM_ and CD8^+^ T_EM_ cells were higher in GL261-Red-FLuc TB brains (**Fig. S10**). At day 7, CD4⁺ T_CM_ cell abundance in GL261-Red-FLuc TB brains (2.5 ± 1.9% of CD4^+^ cells) was 7.3-fold and 3.9-fold higher than parental GL261 (0.35 ± 0.44% of CD4^+^ cells; *p* = 0.01) and GL261-NLuc (0.65 ± 0.54% of CD4^+^ cells; *p* = 0.03) TB brains, respectively (**Fig. 4C**); by day 21, CD8⁺ T_EM_ cell abundance (50.2 ± 9.0% of CD8^+^ cells) increased by 1.5-fold (34.0 ± 7.6% of CD8^+^ cells; *p* = 0.003) and 1.6-fold (30.5 ± 3.5% of CD8^+^ cells; *p* < 0.001), respectively (**Fig. 4D**). GL261-Red-FLuc TB brains were also characterized by greater granzyme B (GZB) expression on day 21, where GZB-expressing total CD8^+^ T cells (3.4 ± 1.2% of CD8^+^ cells) increased 2.4-fold compared to parental TB brains (1.4 ± 0.7% of CD8^+^ cells; *p* = 0.02; **Fig. 4E**). Not surprisingly, PCA with all the described T cell markers clustered GL261-WT and GL261-NLuc together and separated them from GL261-Red-FLuc (**Fig. 4F**), confirming similar T cell activation landscape in parental GL261 and GL261-NLuc TB brains and distinct from GL261-Red-FLuc.

### GL261-NLuc and GL261-WT show sex-independent clustering similarity in exploratory analyses

We also repeated the study using female mice. As in males, GL261-Red-FLuc tumors implanted into female mouse brains regressed (*data not shown*). The immune composition in tumor-implanted TB brains changed between days 7 and 21 p.i. across all groups (**Fig. 5A**; **Fig. S11A**), while remaining stable in contralateral hemispheres (*data not shown*). At day 21 the immune cell composition in TB brains—predominantly MDM, NK, and T cells—was also comparable between males and females (**Fig. 3A**; **Fig. 5A**). The abundance of other immune cell subsets within implanted TB brains were also comparable between males and females (**Fig. 3B; Fig. S11A**).

**Fig. 5.**
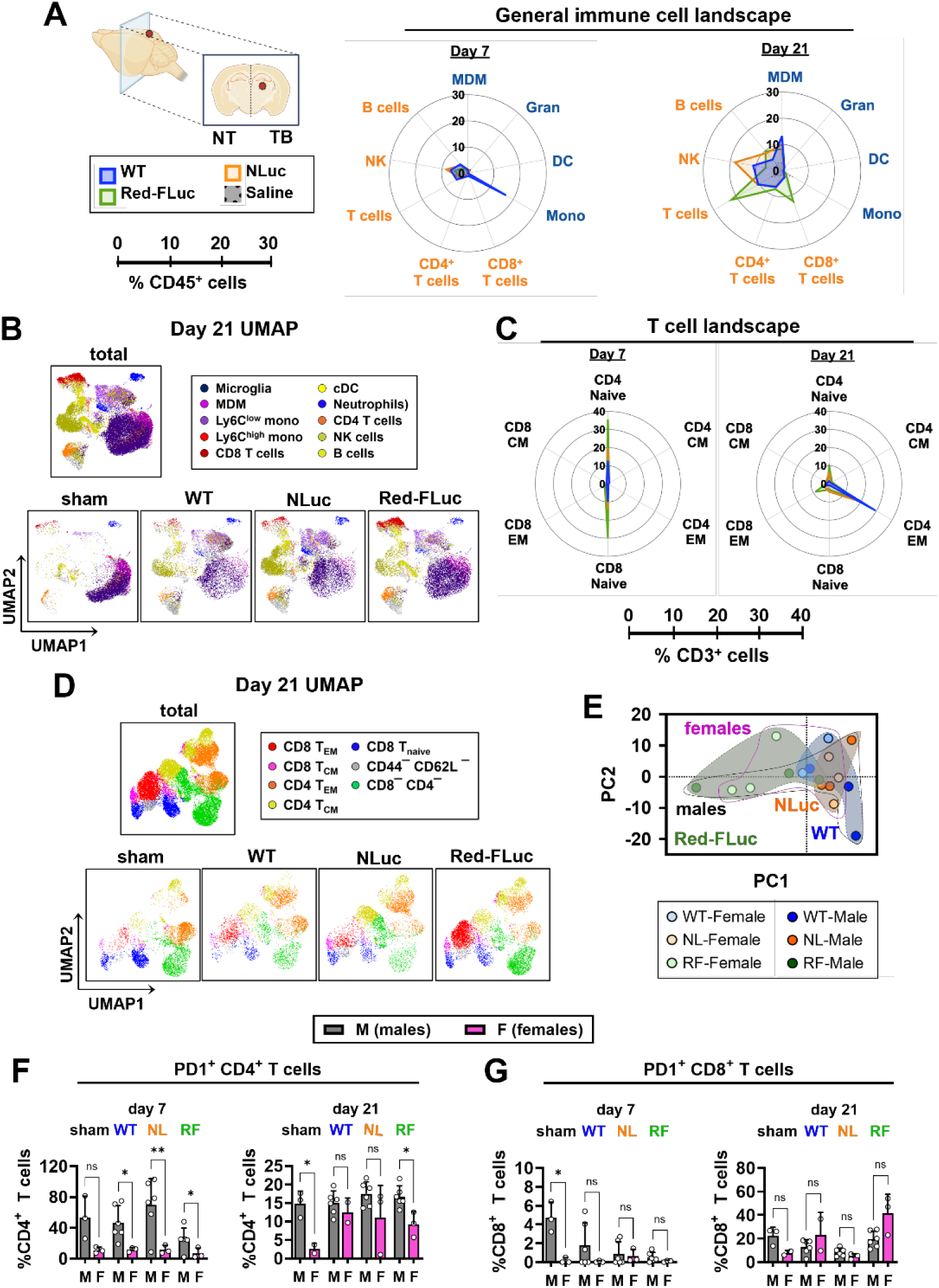
GL261–NLuc tumors retain immune and T cell landscapes comparable to the parental GL261-WT tumors in exploratory male and female cohorts. (**A**) Radar plots showing immune cell populations in tumor-bearing (TB) brains as percentages of CD45+ cells at days 7 and 21 post-implantation of either parental GL261 (*WT*), GL261-NLuc (*NLuc*, *NL*), GL261-Red-FLuc (*Red-FLuc, RF*) cells, or saline (*sham*). (**B**) UMAP visualization of immune cell profiles at day 21 with cell types assigned using manual gating in Fig. S2. (**C**) Radar plots showing T cell populations as percentages of CD3+ cells at days 7 and 21. (**D**) UMAP visualization of T cell profiles at day 21. (**E**) Principal component analysis of immune and T cell composition in male and female TB brains from WT, NL, and RF groups. (**F-G**) PD1 expression on CD4+ T cells (**F**) and CD8+ T cells (**G**) in TB brains. Data are mean ± SD from 6 male and 3 female mice per group; sex differences were analyzed by Welch’s t test. \**p* < 0.05; \*\**p* < 0.005; \*\*\**p* < 0.001; *ns*, not significant. *Gran*, granulocytes; *MDM*, monocyte-derived macrophages; *DC*, conventional dendritic cells; *Mono*, monocytes; T_CM_, central memory T cells; T_EM_, effector memory T.

UMAP visualization of manually gated FCM data from female TB brains at day 21 revealed similar immune landscapes between parental and GL261-NLuc, distinct from GL261-Red-FLuc which contained overtly increased abundance of microglia and CD4^+^ and CD8^+^ T cells (**Fig. 5B**). The amounts of microglia (32.9 ± 3.5% of CD45^+^ cells), CD4⁺ T cells (7.5 ± 0.6% of CD45^+^ cells), and CD8⁺ T cells (15.7 ± 1.3% of CD45^+^ cells) in GL261-Red-FLuc TB brains were increased by 1.7-fold (19.5 ± 0.6% of CD45^+^ cells; *p* = 0.04), 2.0-fold (3.8 ± 0.7% of CD45^+^ cells; *p* = 0.005), and 6.5-fold (2.4 ± 1.4% of CD45^+^ cells; *p* < 0.001), respectively, relative to parental TB brains (**Fig. S12**). These FCM observations were further validated using unsupervised FlowSOM analysis (**Fig. S13**) with observed patterns consistent with male TB brains (**Fig. S6**).

As in males, dynamics in CD4⁺ and CD8⁺ T cell populations in female TB brains similarly exhibited a shift from T_naive_ to T_CM_ and T_EM_ states (**Fig. 4A**; **Fig. 5C**). The T cell landscapes in GL261-NLuc and parental TB brains did not differ from each other but differed from GL261-Red-FLuc (**Fig. S14**). On day 7, GL261-Red-FLuc TB brains contained 2.8-fold higher CD4⁺ T_naive_ (35.6 ± 2.5% of CD4^+^) and 3.0-fold higher CD8⁺ T_naive_ (29.9 ± 2.2% of CD8^+^) cell abundance than in parental GL261 TB brains (CD4⁺ T_naive_: 12.7 ± 8.1% of CD4^+^; *p* = 0.003; CD8⁺ T_naive_: 9.9 ± 8.2% of CD8^+^; *p* = 0.01); by day 21, CD8⁺ T_EM_ cells (8.8 ± 4.1% of CD8^+^) in GL261-Red-FLuc TB brains were increased 4.0-fold relative to parental GL261 (2.2 ± 0.2% of CD8^+^; *p* = 0.03) (**Fig. S14**). These findings were corroborated by UMAP analysis in manually gated FCM data, which highlights the overt increase in CD8^+^ T_EM_ in GL261-Red-FLuc TB brains compared to other groups (**Fig. 5D; Fig. S11C**).

Most importantly, PCA with combined general immune and T cell compartments clustered GL261-WT and GL261-NLuc together, distinct from GL261-Red-FLuc, without overt segregation by sex (**Fig. 5E**). The primary sex-associated difference discernible from the markers we evaluated was the higher frequency of PD1⁺ expression in CD4⁺ T cells in male *vs.* female TB brains (**Fig. 5F**). On day 7, the frequency of PD1 expression in male GL261-WT (46.7 ± 22.2% CD4^+^), GL261-NLuc (69.9 ± 34.2% CD4^+^), or GL261-Red-FLuc (25.2 ± 14.5% CD4^+^) TB brains was 4.0-fold, 6.0-fold, and 3.6-fold higher than in female brain counterparts (WT: 11.6 ± 2.9% CD4^+^, *p* = 0.01; NLuc: 11.8 ± 5.4% CD4^+^, *p* = 0.008; Red-FLuc: 7.0 ± 6.9% CD4^+^, *p* = 0.04) (**Fig. 5F**). This difference persisted to day 21 in GL261-Red-FLuc brains (16.7% in males *vs.* 9.2% CD4^+^ in females, *p* = 0.04) (**Fig. 5F**). A similar trend in PD1⁺ expression frequency on CD8⁺ T cells was only observed in sham implanted brains on day 7 (**Fig. 5G**). Interestingly in male TB brains, PD1 expression on CD4⁺ T cells decreased between days 7 and 21 but increased on CD8⁺ T cells; however, PD1 expression remained stable in females— except in CD8^+^ T cells from GL261-Red-FLuc brains (**Fig 5F-5G**; **Fig. S15**).

### GL261-NLuc facilitates longitudinal monitoring without altering response to ZIKV-LAV treatment

GL261-WT, GL261-NLuc, and GL261-Red-FLuc cells were infected with Zika virus live-attenuated vaccines (ZIKV-LAV) repurposed as oncolytics for human GBM (*30, 31*). *In vitro* infectivity assays revealed similar infection susceptibility between GL261-NLuc cells and parental cells. Compared to mock-infected parental GL261 cells, viability at day 2 post-infection was most affected by ZIKV-LAV DN-1 (35% reduction, *p* = 0.005) and least affected by ZIKV-LAV DN-2 (9% reduction, *p* = 0.24); this trend was mirrored in GL261-NLuc cells (**Fig. 6A**). In comparison, GL261-Red-FLuc cell viability was significantly diminished by all viruses: 32%, 80%, and 54% reduction with ZIKV HPF (*p* = 0.03), ZIKV-LAV DN-1 (*p* = 0.008), and DN-2 (*p* = 0.002) infection, respectively (**Fig. 6A**). These observations were reproduced at day 5 post-infection (**Fig. 6B**).

**Fig. 6.**
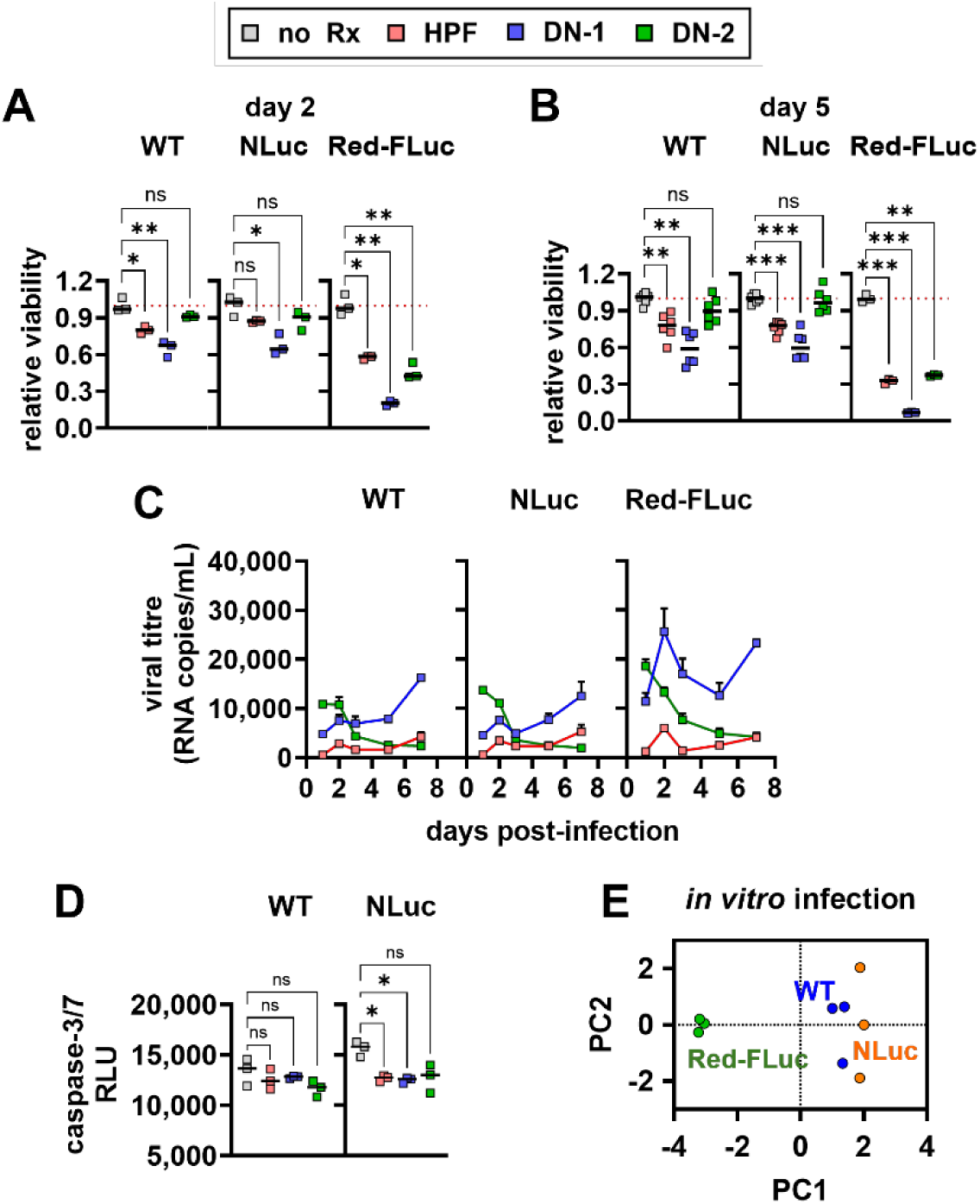
NLuc expression in GL261 glioma cells does not alter susceptibility to ZIKV-LAV infectivity, replication, or cytotoxicity. (**A**-**B**) Viability of parental GL261 (*WT*), GL261-NLuc (*NLuc*), and GL261-Red-FLuc (*Red-FLuc*) cells after infection with Zika virus (*ZIKV*) parental strain HPF or ZIKV live-attenuated vaccine (ZIKV-LAV) strains DN-1 or DN-2 at a multiplicity of infection of 1 for duration of 2 days (**A**); or 5 days (**B**). (**C**) Viral replication kinetics in infected cells measured by qRT-PCR from days 1 to 7 post-infection. (**D**) Apoptosis measured using the caspase-3/7-Glo assay. (**E**) Principal component analysis of *in vitro* infectivity and cytotoxicity profiles across cell lines. Data in **A**, **B**, and **D** were reported as mean ± SD and analyzed by Welch’s ANOVA with Dunnett’s correction. \**p* < 0.05; \*\**p* < 0.005; \*\*\**p* < 0.001; *ns*, not significant.

Viral titers over a 7-day infection period were comparable between parental GL261 and GL261-NLuc cells. ZIKV-LAV DN-1 titers peaked on day 7 (1.6×10^4^ copies/mL *vs.* 1.2×10^4^ copies/mL) and DN-2 titers peaked on day 1 (1.1×10^4^ copies/mL *vs.* 1.4×10^4^ copies/mL), while the respective titers in GL261-Red-FLuc cells were comparable (1.6×10^4^ copies/mL for DN-1, and 1.8×10^4^ copies/mL for DN-2) (**Fig. 6C**). However, ZIKV-LAV DN-1 titers in GL261-Red-FLuc cells peaked on day 3 (2.6×10^4^ copies/mL), with 3-fold higher values than wild type and GL261-NLuc on the same day (**Fig. 6C**). Compared to parental and GL261-NLuc cells, GL261-Red-FLuc cells were more permissive to infection and supported replication of DN-1 and DN-2 as early as 2 days post-infection (**Fig. 6C**).

Despite minimal virus-induced cell death in GL261-WT and GL261-NLuc cells (**Fig. 6A-B**), apoptosis was evident from caspase-3/7 detection in cell lysates (**Fig. 6D**).

Based on these virological assessments, PCA independently clustered GL261-NLuc with the parental cells and separated them from GL261-Red-FLuc (**Fig. 6E**), indicating that NLuc expression does not materially alter the susceptibility of GL261 cells to ZIKV-LAV infection *in vitro*.

We next evaluated the *in vivo* performance of GL261-NLuc in an oncolytic virotherapy setting by implanting immunocompetent mice with cells pre-infected with ZIKV-LAV. Survival curves of mice implanted with mock-infected parental GL261 cells (OS_m_ = 22.5 days) were comparable to mice implanted with cells pre-infected with virus (21.5 days *vs.* 26.5 days *vs.* 21.5 days in DN-1, DN-2, and HPF, respectively; *p* = ns) (**Fig. 7A**). Moreover, ZIKV-LAV pre-treatment of cells prior to implantation did not reduce the clinical burden of disease, as shown by body weight records (**Fig. S16**), and postmortem tumor sizes (**Fig. S17**). Despite absence of overt survival benefit, 50% of mice implanted with parental GL261 cells pre-infected with ZIKV-LAV DN-2 was still alive at 26 days p.i.—*i.e.,* when all mock-treated mice succumbed to disease (**Fig. 7A**).

**Fig. 7.**
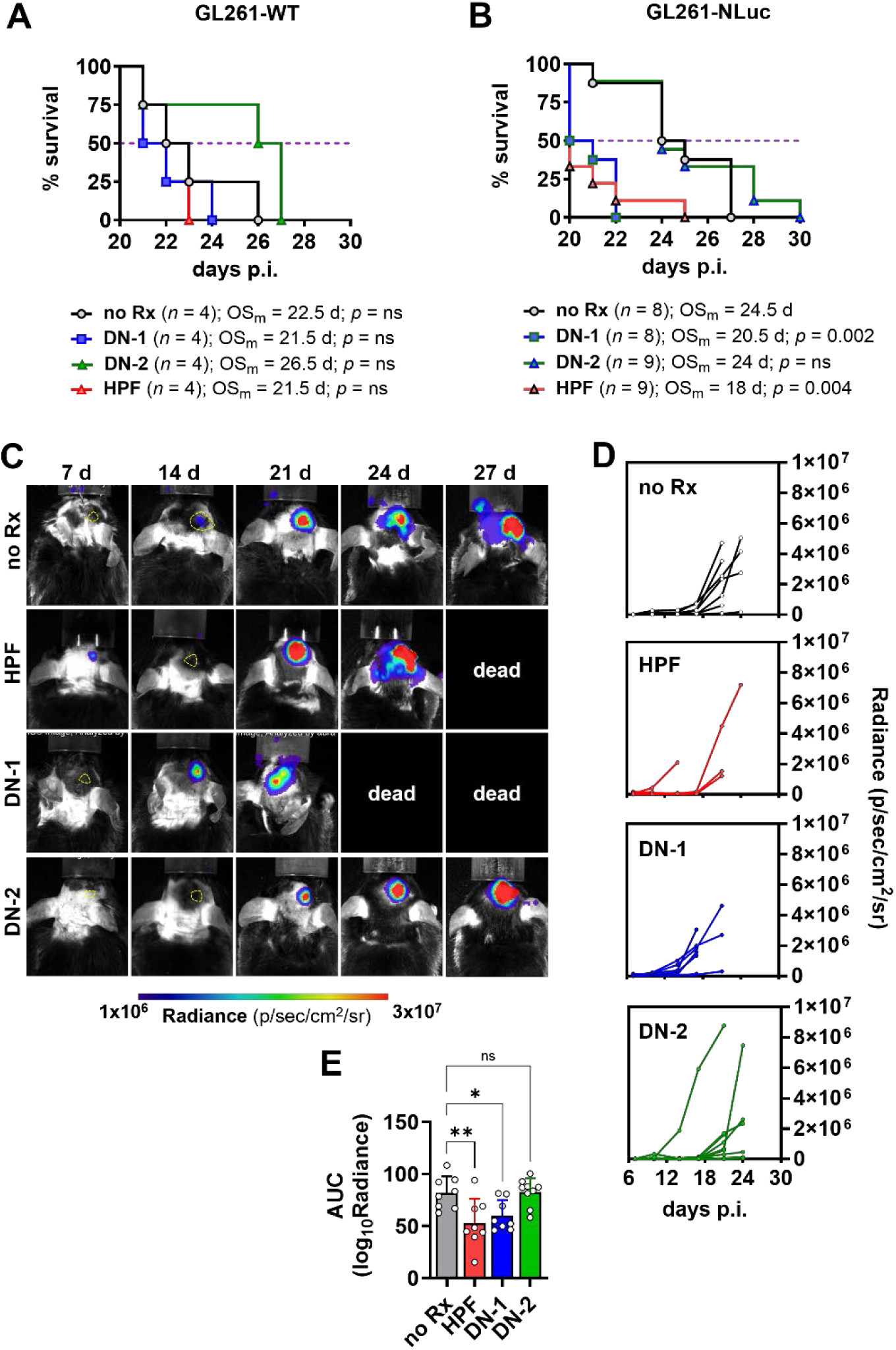
Bioluminescence imaging of tumors in mice reveals temporary control of glioma growth following ZIKV-LAV infection. (**A**-**B**) Kaplan–Meier survival of mice implanted with GL261-WT (**A**); or GL261-NLuc (**B**) cells infected with either Zika virus parental strain (*HPF*) or ZIKV live-attenuated vaccine (*DN-1* or *DN-2*) at a multiplicity of infection of 1; or saline (*no Rx*). (**C**-**E**) Representative images of longitudinal *in vivo* bioluminescence imaging (BLI) of tumor-bearing mice (**C**). (**D**) Quantification of luminescence signals over time in mice implanted with GL261-NLuc cells pre-exposed to HPF (*n* = 9), DN-1 (*n* = 8), DN-2 (*n* = 9), or mock treatment (*n* = 8). (**E**) Area under the curve (AUC) analysis of log-transformed BLI data. Median survival (OS_m_) relative to mock-treatment in **A-B** were compared using Mantel-Cox test with Holm-Šídák correction. **E** is shown as mean ± SD and analyzed by one-way ANOVA with Dunnett’s correction. \**p* < 0.05; \*\**p* < 0.005; \*\*\**p* < 0.001; *ns*, not significant. Tumor regions in **C** are outlined with yellow dashed lines.

On the other hand, mice implanted with mock-infected GL261-NLuc or pre-treated with ZIKV-LAV DN-2 exhibited comparable survival (OS_m_ 24.5 *vs.* 24 days) and disease burden (**Fig. 7B; Fig. S16, Fig. S17**). Despite this, 22% of mice implanted with ZIKV-LAV DN-2-pre-treated cells survived beyond day 27 p.i.—*i.e.,* when all the mock-treated mice succumbed (**Fig. 7B**). In contrast, mice implanted with GL261-NLuc pre-exposed to ZIKV HPF or ZIKV-LAV DN-1 exhibited shorter survival time compared to mock treatment (18 days, *p* = 0.004; and 20.5 days, *p* = 0.002, respectively) (**Fig. 7B**).

Crucially, longitudinal BLI of mice implanted with GL261-NLuc cells pre-exposed to ZIKV-LAV DN-2 revealed a temporal delay of 2-3 days in exponential increase of radiance compared to those implanted with either mock-treated cells or cells pre-exposed to ZIKV HPF (**Fig. 7C-D**). AUC analysis of longitudinal imaging data revealed pre-treatment of GL261-NLuc cells with ZIKV HPF and ZIKV-LAV DN-1 reduced tumor growth kinetics by 42.9% (± 22.0%, *p* = 0.005) and 26.6% (± 18.2%, *p* = 0.04), respectively, compared to mock-infection treatment despite the lack of survival benefit and even earlier death in these mice (**Fig. 7E**).

## Discussion

We engineered GL261 mouse glioma cells to stably express NLuc, benchmarked the *in vitro* properties of GL261-NLuc with parental GL261 cells, and established an immunocompetent, syngeneic orthotopic brain tumor model compatible with non-invasive longitudinal BLI. This work was prompted by accumulating evidence that transgenic reporters, including luciferases, can alter tumor growth kinetics and immunogenicity in immunocompetent hosts (*20–22, 32, 33*). Several GL261 reporter lines expressing firefly luciferase variants, including Red-FLuc (*34*), have been shown to elicit anti-tumor immune responses, increased inflammatory signaling, and altered T cell infiltration, thereby compromising the fidelity of GBM models (*19, 23, 35*). We therefore systematically evaluated whether NLuc could provide a sensitive imaging platform while preserving the biological and immunological characteristics of the parental GL261 model.

*In vitro*, GL261-NLuc retained the morphology, growth kinetics, and overall proliferative behavior of parental GL261, consistent with previous reports showing that luciferase expression does not necessarily alter GL261 growth under standard culture conditions (*17, 36*). Although clonogenic growth was modestly reduced at low seeding density, this likely reflects altered paracrine dependence rather than intrinsic proliferative defects (*37, 38*). Importantly, NLuc generated over 100-fold greater signal intensity than Red-FLuc, consistent with its high catalytic efficiency (*39*), providing substantially improved sensitivity for detecting tumor growth and growth-associated changes over time.

*In vivo*, GL261-NLuc formed lethal intracranial tumors that exhibited histopathology and molecular features closely resembling those in parental GL261 TB brains. Meanwhile, GL261-Red-FLuc-derived tumors regressed and corroborated previous reports of poor engraftment and reporter-driven immunogenicity accompanied by greater immune-cell infiltration and exacerbated inflammatory signaling in C57BL/6 mice (*19, 23*). Unlike Red-FLuc, NLuc expression permitted sustained tumor growth in immunocompetent hosts while maintaining the major biological and pathological characteristics of the parental model.

A central finding of this study was that implantation of GL261-NLuc cells largely preserved the immunosuppressive tumor microenvironment characteristic of parental GL261 tumors. Cytokine profiling, flow cytometry, and multivariate analyses of TB brains consistently clustered GL261-NLuc with GL261-WT and separated both from GL261-Red-FLuc tumors. GL261-Red-FLuc cell-implanted TB brains were associated with enhanced cytotoxic T-cell recruitment, elevated inflammatory signaling, and broad immune activation, suggesting a reporter-driven immune response that substantially altered tumor biology. Collectively, these findings indicated that NLuc causes considerably less immunological perturbation than Red-FLuc while largely preserving immune microenvironment fidelity.

To strengthen our FCM-based immune landscape analyses, we employed both conventional manual gating and unsupervised FlowSOM clustering. Whilst our manual gating strategy ensured the differentiation of immune cell types according to our pre-defined preferences, FlowSOM facilitated unbiased clustering that takes into consideration the variability in expression levels of all stained biomarkers. The high concordance between these independent analytical approaches supported the robustness of the observed immune phenotypes and reduced the likelihood that the findings were simply artefacts of a particular clustering strategy.

Exploratory sex-stratified analyses similarly suggested that the overall parity between GL261-NLuc and parental GL261 was maintained in both male and female mice, and PCA could not segregate the groups by sex. However, the small female cohort in this study precludes formal conclusions regarding sex equivalence. Across the measured immune parameters, higher PD1 expression on tumor-infiltrating T cells in males *vs.* females was the primary sex-differentiating factor, consistent with growing evidence that immune checkpoint regulation can be influenced by sex (*40–42*). These observations reinforce the importance of considering sex as a biological variable, particularly in studies investigating PD1/PD-L1-directed immunotherapies.

We also investigated whether GL261-NLuc retained the response characteristics of parental GL261 under biological perturbation using ZIKV-LAV treatment (*30, 31*). *In vitro*, GL261-NLuc and GL261-WT displayed comparable susceptibility to viral infection, viral replication, and virus-associated growth inhibition; in contrast, GL261-Red-FLuc cells exhibited heightened permissiveness to virus infection. Of the two ZIKV-LAV strains, DN-1 exhibited a potent inhibitory effect on glioma growth *in vitro*, consistent with findings from treatment of human GBM cell lines (*31*). *In vivo*, longitudinal BLI revealed treatment-associated changes in tumor growth kinetics that were not readily apparent from survival analyses alone. Pre-exposure of GL261-NLuc cells to ZIKV-LAV DN-2 delayed tumor outgrowth and reduced overall signal accumulation in mice. Pre-exposure to ZIKV HPF and ZIKV-LAV DN-1, likewise, slowed tumor growth despite causing earlier death. These findings highlight an important advantage of NLuc-based longitudinal imaging: the ability to sensitively quantify dynamic changes in tumor progression that may not be captured by conventional endpoint measures. Rather than establishing the therapeutic efficacy of ZIKV-LAV, these experiments demonstrated that GL261-NLuc retains the response characteristics of parental GL261 under viral perturbation and provides a useful platform for monitoring treatment-associated changes in tumor growth.

Unexpectedly, ZIKV DN-1 pre-exposure of parental and GL261-NLuc cells prior to mouse brain implantation resulted in earlier mortality despite showing potent anti-glioma activity *in vitro* and previously reported attenuation in immunocompromised mice (*30*). One possible explanation is that implantation of virus-exposed tumor cells directly into the brain facilitated local viral infection of neural tissue. Additionally, differences in host age and immune status between the present model and previous vaccine studies may have increased susceptibility to neuroinflammatory complications.

Several limitations are worth noting in this study. First, although GL261-NLuc retained key tumor and immune features of the parental model, we have only shown this using a single homogeneous glioma line, which does not fully capture the molecular heterogeneity of human GBM. Second, although NLuc expression was associated with markedly less immune perturbation than Red-FLuc, this does not mean NLuc-driven immune effects were entirely absent. Larger biomarker panels in FCM would be necessary for deeper immune profiling and more accurate distinction of cell types in the tumor microenvironment. Third, the reliance on BLI, while enabling high-throughput longitudinal monitoring, provided limited spatial resolution and might not fully reflect tumor architecture or molecular profiles compared to modalities such as MRI or PET, respectively. Substrate delivery and signal attenuation in deep brain tissue might also introduce variability in signal quantification. Future studies could leverage novel NLuc substrates that cross the blood-brain barrier more easily and were specifically developed for brain imaging (*43, 44*). Fourth, from a statistical perspective, the study may be underpowered to detect subtle differences in some immune or therapeutic endpoints, particularly in subgroup analyses stratified by sex. Sample sizes were sufficient to resolve major phenotypic differences between models but could not detect smaller effect sizes. Finally, therapeutic evaluation was restricted to a single oncolytic platform (ZIKV-LAV), and broader validation across additional therapeutic modalities would be necessary to establish the generalizability of these findings.

In conclusion, stable NLuc expression enables sensitive longitudinal imaging of orthotopic gliomas while largely preserving the biological, molecular, and immunological characteristics of the widely used GL261 model. Compared with Red-FLuc, NLuc was associated with markedly less immune perturbation, maintained tumor progression and therapeutic response profiles, and preserved key features of the parental tumor microenvironment. These properties address a longstanding challenge in preclinical glioblastoma research: the need for longitudinal, non-invasive tumor monitoring with minimal reporter-associated immune artefacts. As immunotherapies, cancer vaccines, and oncolytic virotherapies increasingly depend on accurate assessment of tumor–immune interactions, the availability of a high-sensitivity reporter model with comparatively limited immunologic perturbation will improve the reliability and interpretability of longitudinal studies in immunocompetent hosts. Beyond glioblastoma, these findings highlight the importance of evaluating reporter immunogenicity as a critical determinant of model fidelity and provide a framework for the development of next-generation imaging-compatible cancer models.

## Materials and Methods

### Experimental design

We investigated the phenotypic changes induced by imaging reporter expression in GL261 murine glioma cells both *in vitro* and *in vivo*. *In vitro* growth kinetics of parental GL261 cells, as well as cells expressing either NLuc or Red-FLuc, were investigated by comparing cell morphology and cell doubling times under various cell seeding densities. Lastly, the stable expression and brightness of NLuc in GL261 cells were also compared. Studies were repeated in three independent experiments with 6 samples each.

For studies on survival, gross pathology, and histopathology, male C57BL/6N mice were intracranially implanted with either parental cells (*n* = 5), GL261-NLuc (*n* = 8) or GL261-Red-FLuc (*n* = 10) and observed for 30 days. *In vivo* BLI was performed in mice implanted with reporter-expressing cells every 7 days. Brains harvested from mice that succumbed on day 21 were used in gross pathology, histopathology, and protein expression (cytokine and chemokine) assays.

Studies on immune landscape profiling were conducted using 2 cohorts of male mice and 1 cohort of female mice. Across two cohorts, 12 male mice each were intracranially implanted with either GL261-WT, GL261-NLuc, GL261-Red-FLuc, or saline (total of 48 male mice). Six male mice from each group were euthanized on day 7, and the remaining 6 mice on day 21 p.i. Similarly, 6 female mice each were intracranially implanted with either GL261-WT, GL261-NLuc, GL261-Red-FLuc, or saline (total 24 mice). Three female mice from each group were euthanized on day 7, and the remaining 3 mice on day 21 p.i.

### Cells

Unmodified GL261 (GL261-WT) cells were obtained from the National Cancer Institute (NCI), while GL261 cells expressing the red-shifted firefly luciferase (Bioware^®^ Brite GL261-Red FLuc) were commercially sourced (PerkinElmer [now Revvity], USA). GL261 cells expressing nanoluciferase (GL261-NLuc) were generated through lentiviral transduction (ATUM Bio, USA).

### Production of recombinant lentivirus

HEK293T cells were transiently transfected with third-generation lentiviral vector DNA (Addgene) and the NLuc lentiviral expression vector (ATUM Bio, USA) using FuGENE 4K transfection reagent (Promega) in DMEM (Gibco, Thermo Fisher Scientific) supplemented with 10 % fetal calf serum (FCS; HyClone). Media was replaced with fresh DMEM (10% Fetal Calf Serum, FCS; 2 mM caffeine; pH 6.3) at 16-18 h post-transfection. Lentivirus culture supernatant was collected 24 h and 48 h post-transfection, filtered (0.45 µm), and concentrated using Lenti-X Concentrator (Clontech, Takara Bio, Japan). The resulting viral pellet was resuspended in serum-free DMEM and used freshly for transduction of target cells, or stored at −80 °C.

### Generation of GL261-NLuc cells

GL261-WT cells cultured in DMEM (10 % FCS) were transduced with concentrated virus and 8 µg/mL hexadimethrine bromide (polybrene, Sigma-Aldrich) and cultured overnight. The viral supernatant was replaced with fresh DMEM (10% FCS) at 18 h post-transduction, and successfully transduced cells were selected with puromycin (5 µg/mL, Invivogen).

### In vitro cell characterizations

Live cell imaging and growth kinetics were established using Incucyte S3 (Sartorius, Germany). Briefly, 3,000 cells were seeded in 96-well plates (9,645 cells / cm^2^), and live-cell image capture was performed every 6 h over 5 days. Cell confluence (from 0-100%) was evaluated longitudinally using a cell mask prepared for GL261 cells, and data were fitted into a logistic growth curve with set constraints (Y_m_ = 100) in Prism v.10.1 software (GraphPad, USA). Cell doubling time (T*_d_*) was calculated using the following formula: 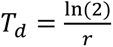, where *r* represents the maximum per capita growth rate at the inflection point, calculated from the logistic equation.

Uncontrolled cell proliferation was evaluated using colony formation assays as described previously (*31*). Briefly, 2,000 cells were seeded in 24-well plates (1,052 cells / cm^2^) and incubated in DMEM (10% FCS) for 2 weeks with fresh media replacement every 3-4 days. Upon colony formation, cells were fixed in 4% paraformaldehyde, stained with 4% alcoholic crystal violet, and imaged with the Gel Count^TM^ automated colony counter (Scintica, USA). Crystal violet was subsequently extracted from fixed cells with mild detergent (2% SDS), and the absorbance at 570 nm was measured using a plate reader (Tecan, Switzerland).

Expression of NLuc in stably transduced cells was evaluated by *in vitro* bioluminescence imaging (BLI) assays. Briefly, GL261-NLuc cells were seeded overnight at two-fold serial dilution from 50,000 to 3,125 cells per well. At 18 h to 24 h post-seeding, cells were supplemented with fluorofurimazine (Ffz; nanoGLO^®^, Cat. No. N1110; Promega, USA) and subjected to BLI using the Sapphire Biomolecular Imager (Azure Biosystems, USA). Similarly, Red-FLuc luminescence was measured from GL261-Red-FLuc cells using D-luciferin (Gold Biotechnology, USA). Luminescence signals were quantified using ImageJ (National Institutes of Health (NIH), USA).

Stable NLuc expression was also evaluated by standard Western blot at various stages of cell passaging using anti-NLuc antibody (Cat. No. N7000; Promega, USA). Protein expression was quantified by densitometry using ImageJ (NIH, USA) and normalized to expression of the housekeeping protein β-actin (Cat. No. 66009; Proteintech, Singapore).

### Ethics statement

All animal experiments were conducted with approval from the Institutional Animal Care and Use Committee (IACUC) of Duke-NUS Medical School and SingHealth (IACUC approval no. 2019/SHS/1533).

### Orthotopic mouse glioma model

6- to 7-week-old C57BL/6N mice (adult male and female) were purchased from a local colony (InVivos, Singapore) and acclimated for 3-4 days at Duke-NUS vivarium prior to experiments. WT and GL261 cells expressing imaging reporters were implanted orthotopically into mouse brains following published protocols with modifications (*16, 45*). Briefly, mice were anesthetized with a cocktail of ketamine (100 mg/kg) and xylazine (20 mg/kg) by intraperitoneal (i.p.) injection prior to the procedure. Heads were depilated using hypoallergenic shaving cream one day prior to the tumor implantation to expose the scalp. Anesthetized mice were mounted onto a stereotaxic frame with the head secured by ear bars (Cat. #900-0068-516, 900-0068-605; RWD Life Sciences, USA), and the scalps were disinfected with betadine solution. A small (∼ 1 cm), shallow incision was made on the scalp above the right brain hemisphere—where the cells would be implanted to expose the skull. A burr hole was subsequently created through the skull using an electric microdrill (Cat. #78001; RWD Life Sciences, USA) at the following coordinates relative to the bregma: (AP = −2 mm; ML = 2 mm; DV = −3 mm). GL261-WT, GL261-NLuc, or GL261-Red-FLuc cells (2 ◻ 10^4^ cells suspended in 2 μL volume) were micro-injected into the brain with a Hamilton 10 μL syringe with a 26G needle at a rate of 0.5 μL/ min using a syringe pump (Cat. #72-1823; Harvard Apparatus, USA). Following the injection, the burr hole was patched with bone wax, and the scalp was surgically closed with silk sutures. Mice were observed and given subcutaneous daily injections of Baytril® (enrofloxacin; 5 mg/kg) and Meloxicam (5 mg/kg) for 1 week to ameliorate pain and prevent infection.

The mice were housed in individually ventilated cages and provided with food pellets and water *ad libitum.* Post-implantation, animals were observed daily for signs of disease and underwent general health and neurological examinations. Mice were euthanized when moribund, based on these criteria: (1) loss of >20% body weight relative to day 1 post-implantation, (2) persistent neurological deficit manifesting for >48 h, and (3) immobility or severe lethargy (*46*). Animals were euthanized by inhalation of 10% CO_2_ continuously for 5 min. At euthanasia, brains collected for histopathological examinations were fixed in 10% neutral buffered formalin (NBF) for 48 h prior to tissue processing. Tissues collected for cytokine expression profiling were flash-frozen immediately and stored in −80 °C.

### Monitoring tumor growth kinetics

Optical imaging of brains post-implantation was conducted using the IVIS Lumina *in vivo* imaging system (Caliper Life Sciences, USA). Mice implanted with GL261-Red-FLuc were injected i.p. with D-luciferin LUCK^®^ (150 mg/kg; Gold Biotechnology, USA) and subjected to BLI at 10 min post-injection. Mice implanted with GL261-NLuc were injected i.v. with FFz following the manufacturer’s protocol and subjected to BLI at 2 min post-injection, the peak signal intensity based on dynamic imaging studies. IVIS imaging was conducted at the highest stage position closest to the camera; and images were acquired with large binning. Bioluminescence data were analyzed using Aura v.3.2 software (Spectral Instruments Imaging, USA).

### Pathological examination of brain tissues

Upon necropsy, brains were cut in the sagittal midline to separate the tumor-bearing (TB) brain hemisphere from the non-tumor (NT) brain hemisphere for gross pathological examination. Tissues were immediately fixed in 10% neutral buffered formalin (NBF) and equilibrated stepwise in 15% and 30% sucrose prior to gross pathological examination of tumors. Equilibrated brains were embedded in OCT medium, frozen down, and sectioned using a cryotome. Fixed-frozen tissue sections (10 μm) were stained with Harris’ hematoxylin and eosin (H&E) as previously described (*46*).

### Immune profiling of brain tumors and other tissues

Expression of various cytokines in TB brains was determined using the Proteome Profile Mouse XL Cytokine Array (R&D Systems, USA) and following the manufacturer’s protocol. The Western blot hybridized membranes were imaged using Sapphire Biomolecular Imager (Azure Biosystems, USA), and blot intensities were quantified by densitometry using ImageJ (NIH, USA). Three mice were used per group.

Immune cell profiling of TB and NT brains was conducted with flow cytometry (FCM) as previously described (*47, 48*). Three mice per group per time point was used per cohort, and two cohorts of male mice were independently evaluated for data reproducibility. Only one cohort was evaluated for female mice. Harvested tissues were mechanically disaggregated and incubated in digestion buffer containing DNase I and collagenase. Following homogenization, single cells were collected by passing the suspension through a nylon mesh strainer (70 μm). Fat was subsequently removed by centrifugation in Percoll. Cells were labelled with fluorescent antibody cocktails targeting various immune cells and were assayed by flow cytometry (Fortessa, BD Biosciences, USA) and data analysis using FlowJo V10.8.0 (BD Biosciences, USA). Immune cell subsets were identified using a gating strategy as described (*49*) and detected events were either expressed as %CD45^+^ cells or normalized to absolute cell number using CountBright Absolute Counting Beads (ThermoFisher, USA). The following markers were used to identify immune cell subsets: total immune cells (CD45^+^); granulocytes (Ly6G^+^Ly6C^+^CD11b^+^); monocytes (CD11b^+^CD115^+^); microglia (CD11b^+^Ly6G^◻^Ly6c^◻^CD3^◻^); monocyte-derived macrophages (MDM; CD11b^+^Ly6G^◻^Ly6c^+^CD3^◻^); conventional dendritic cells (DC; CD11c^+^MHCII^+^); B cells (CD3^◻^CD19^+^B220^+^MHCII^+^); NK cells (Ly6G^-^Ly6C^-^CD11b^-^CD3^-^CD49b^+^) total T cells (CD19^◻^CD49b^◻^B220^◻^Ly6G^◻^CD3^+^); CD8^+^ T cells (CD19^◻^CD49b^◻^B220^◻^LY6G^◻^CD3^+^CD8^+^), and CD4^+^ T cells (CD19^◻^CD49b^◻^B220^◻^LY6G^◻^CD3^+^CD4^+^). The T cell landscape was evaluated based on the following markers: naïve T cell (CD62L^+^CD44^◻^), central memory T cell (T_CM_; CD62L^+^CD44^+^), and effector memory T cell (T_EM_; CD62L^◻^CD44^+^). In addition, activated CD8^+^ T cells were identified based on granzyme B (GZB) expression, while PD1 expression was measured as an inhibitory/exhaustion-associated marker.

### Visualization of high-dimensional Flow Cytometry Data

FCM data were pre-processed by preparing compensation matrices, biexponential data transformation, and manual gating as described (**Fig. S2**) (using FlowJo). Cell numbers were down-sampled to the same number to avoid unwanted bias, and samples from the same group of mice were concatenated. Uniform manifold approximation and projection (UMAP) was performed with default parameters (neighbors = 15, minimum distance = 0.5, and metric = Euclidean) using the FlowJo UMAP plugin, and the resulting 2D coordinates were plotted to visualize distinct clusters. For validation, unsupervised clustering using the Flow Self-Organizing Map (FlowSOM) plugin (FlowJo) was conducted to define clusters based on marker expression profiles to identify general immune landscape and T cell landscape. Clusters were binned and visualized as an overlay on UMAP plots to compare population distributions across experimental groups.

### Evaluation of GL261 cells using oncolytic viruses

The susceptibility of unmodified (WT) and reporter-engineered (NLuc and Red-FLuc) GL261 cells to infection with Zika virus (ZIKV) live-attenuated vaccine (ZIKV-LAV) strains developed at Duke-NUS (*30*) and subsequently repurposed as oncolytics (*31*) was evaluated. The three strains used included ZIKV parental strain isolated from French Polynesia (H/PF/2013; HPF) and two ZIKV-LAV strains (DN-1 and DN-2), of which DN-2 was demonstrated to be significantly more attenuated both *in vitro* and *in vivo* (*30*).

Briefly, 100,000 cells seeded overnight were inoculated with virus at a multiplicity of infection (MOI) of 1 (*i.e.*, 1 plaque forming unit per cell) in serum-free media for 1 h, followed by replacement with DMEM (2% FCS). For mock-infection, cells were incubated with serum-free media alone. Colony formation assays on infected cells were conducted at 2 and 5 days post-infection (d.p.i.) using alcoholic crystal violet assay described above.

Apoptosis induced by virus infection was measured using caspase-3/7-Glo® luminescence assay at 2 days post-infection (Promega, USA). Lastly, viral replication kinetics in infected cells was determined using qRT-PCR as described previously (*31*). Infected cell supernatants were collected every 12 h, and viral RNA was extracted using QIAamp viral RNA kit (Qiagen, Germany), and viral titers were calculated using a ZIKV RNA standard curve.

### In vivo oncolytic activities in mouse brain tumor implantation models

Mouse tumor models were established as described above using cells pre-infected with ZIKV. Therapeutic intervention of glioma growth kinetics using ZIKV-LAV was evaluated by co-implantation with tumor cells, as described previously (*50, 51*). Overnight-seeded GL261-WT or GL261-NLuc cells were inoculated with either parental ZIKV HPF or ZIKV-LAV strains DN1 or DN2 at 1 MOI. Cells were trypsinized and disaggregated at 24 h post-infection, and 2 ◻ 10^4^ infected cells or sham-treated cells were orthotopically implanted into mouse brains as described above.

Mice were followed up and monitored daily as described above. Mice implanted with infected GL261-NLuc cells were additionally subjected to serial BLI with IVIS Lumina *in vivo* imaging system (Caliper Life Sciences, USA) following intravenous injections with FFz as described above. Once mice were moribund, brains were harvested and processed for immunohistopathological examination as described.

### Statistical analysis and data visualization

Statistical analyses and data visualization were performed with Prism v.10.6.1 Software (GraphPad, USA). Gaussian distribution of continuous data was first determined with Shapiro-Wilk test and QQ plots. Differences among variances between groups following Normal distribution were assessed with Bartlett’s test. Means from two groups of data were compared with either unpaired *t*-test or Welch’s *t*-test for groups with equal variances and unequal variances, respectively. Means from > 2 groups of data with equal variances were compared with one-way ANOVA and Tukey’s correction for multiple comparisons. Similarly, means from > 2 groups of data with unequal variances were compared with Welch’s ANOVA and Dunnett’s correction for multiple comparisons. For data with non-Gaussian distributions, medians were compared with either Mann-Whitney (2 groups) or Kruskal-Wallis (>2 groups) tests with Dunn’s correction for multiple comparisons.

Kaplan-Meier survival curves were compared using Mantel-Cox test with Holm-Šídák correction for multiple comparisons.

Principal component analyses (PCA) in Prism were conducted using “Kaiser rule,” where centered data were converted to eigenvalues, and only principal components (PC) with eigenvalues > 1.0 were selected.

Radar plots and bubble plots were created in Keynote ver. 15.1 (Apple Macintosh, USA), while scientific illustrations and cartoons were created using Biorender©.

## Acknowledgments

**Non-author contributions:** We would like to thank Dr. Masahiro Fukuda (Duke-NUS Neuroscience and Behavioral Disorders (NBD) Programme) for his technical guidance on stereotaxic brain injections. We also acknowledge Dr. Clement Yau (Duke-NUS Emerging Infectious Diseases Programme) for his assistance in producing the batch of ZIKV-LAV strains used in this study.

## Funding

Singapore Health and Biomedical Sciences (HBMS) Industry Alignment Fund Pre-Positioning (IAF-PP) grant H18/01/a0/018 (AMC)

Singapore National Research Foundation (NRF) Grant AG/CIV/GC70-C/NRF/2013/3 (AMC)

Singapore-MIT Alliance for Research and Technology (SMART) Innovation Grant ING-000913 (AMC)

Singapore Ministry of Education Academic Research Tier 1 Grant (FY2022-MOET1-0002)

Duke-NUS Medical School

## Author contributions

Conceptualization: CBLV, RM, AMC

Methodology: CBLV, SHP, RM

Investigation: CBLV, WN, SG, AG, JO

Resources: EEO

Visualization: CBLV, SHP, AMC

Supervision: AMC

Writing—original draft: CBLV

Writing—review & editing: WN, SHP, AMC, RM, EEO

## Data and materials availability

All data are available in the main text or the supplementary materials.

