## Supplemental Files for "Nanoluciferase reporter preserves immunocompetent glioma model fidelity while facilitating longitudinal molecular imaging"

**Fig. S1**

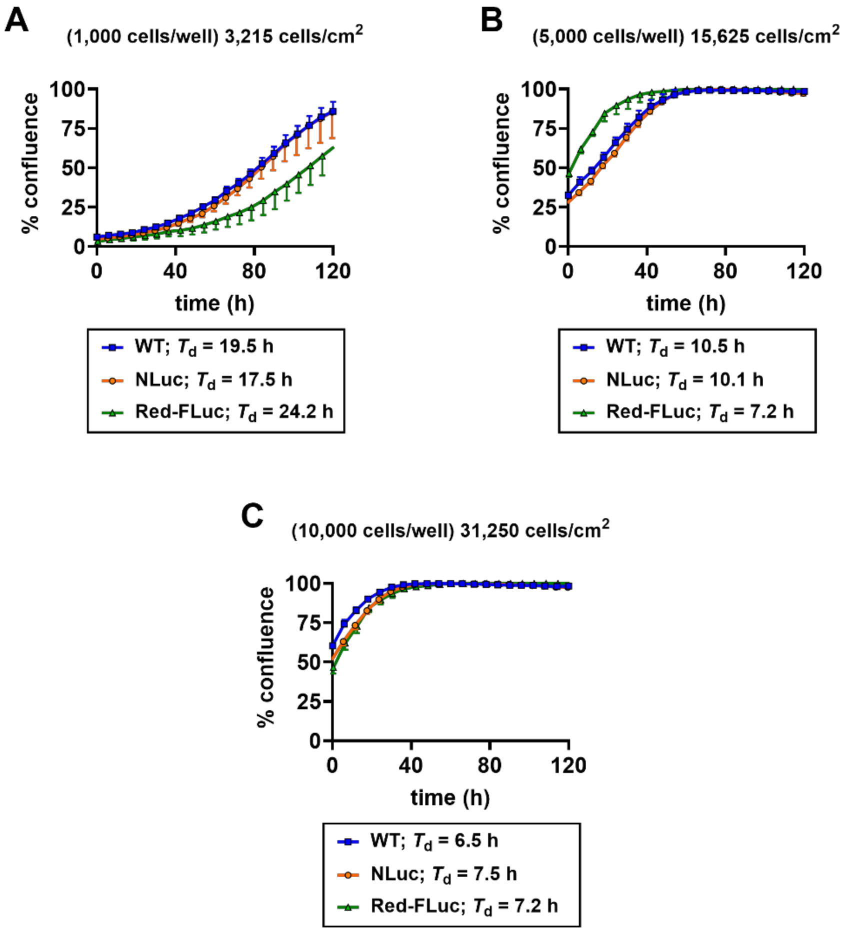


**Fig. S1. Growth curve kinetics at various seeding densities.** Parental (GL261-WT), GL261-NLuc, and GL261-Red-FLuc cells were seeded at various densities: (**A**) 3,215 cells/cm$^2$; (**B**) 15,625 cells/cm$^2$**;** or (**C**) 31,250 cells/cm$^2$. Data are shown as mean ± SD from 6 replicates. T*d*, doubling time.

**Fig. S2**

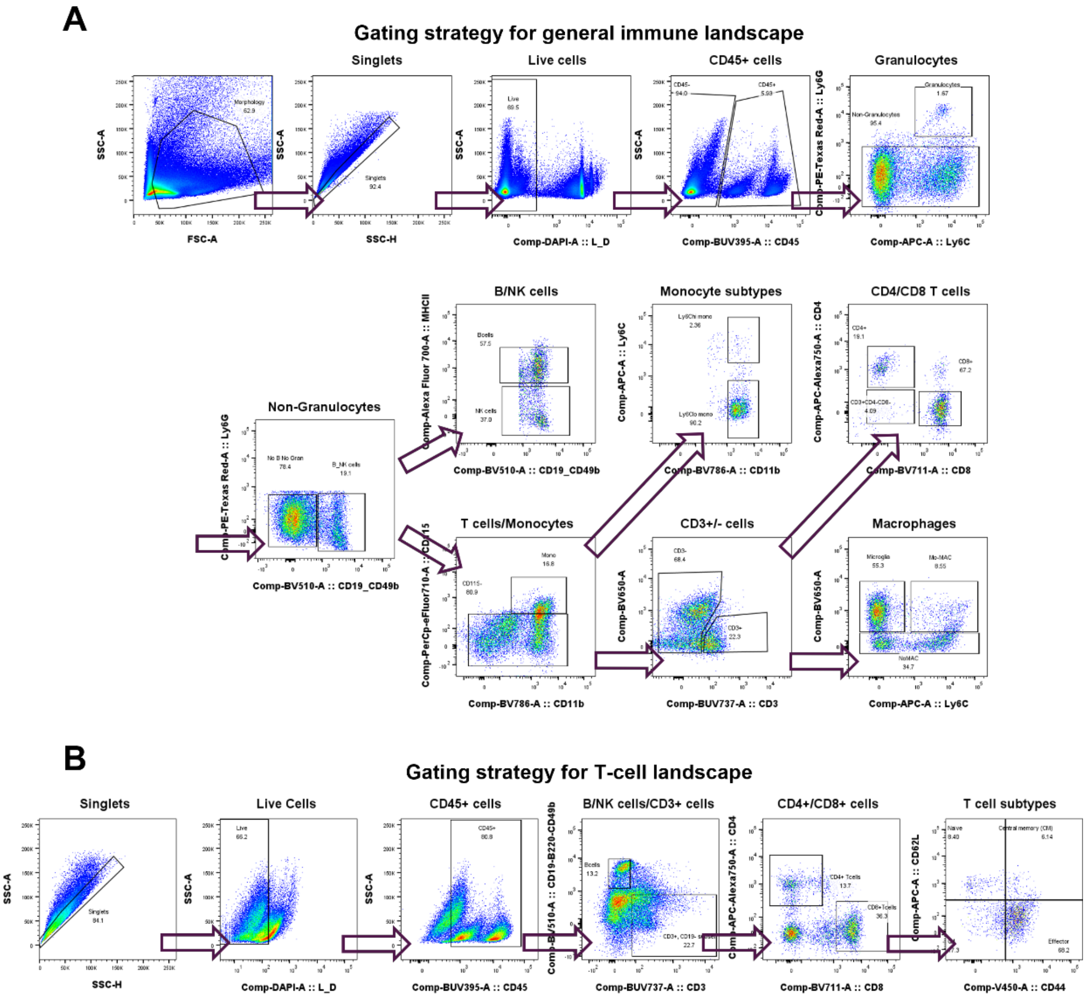


**Fig. S2. Flow cytometry gating strategy for immune and T cell profiling.** Representative gating strategies used to identify (**A**) major immune cell populations and (**B**) T cell subsets in mouse brain tissues.

**Fig. S3**

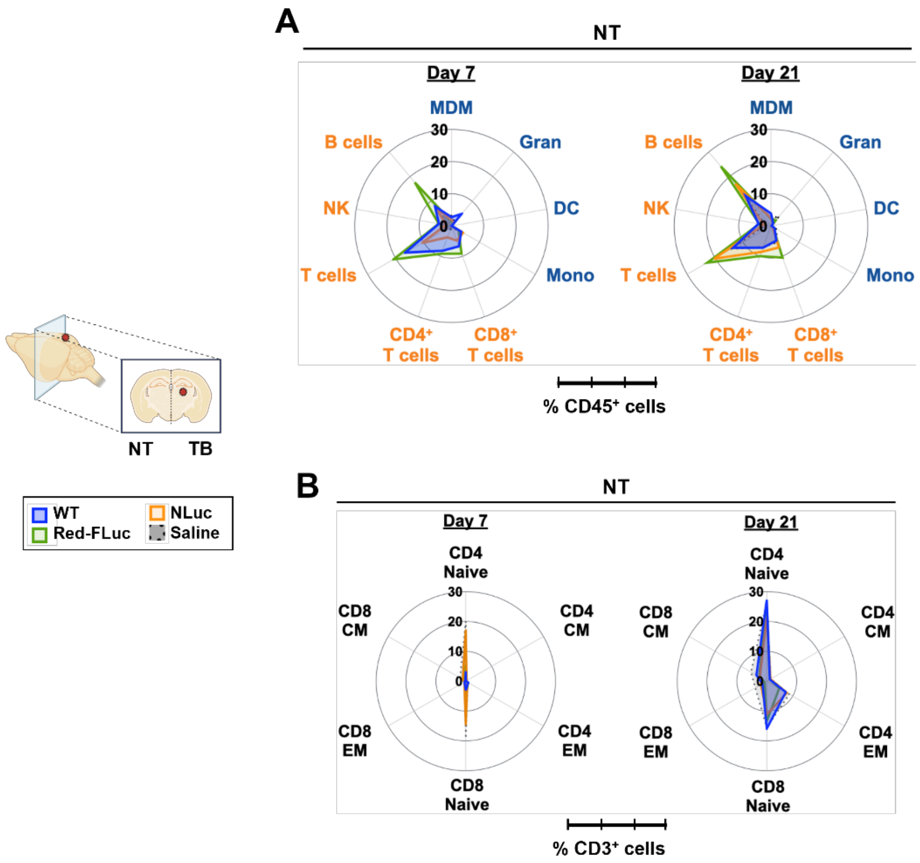


**Fig. S3. Immune and T cell landscapes in contralateral non-tumor-bearing (NT) brain hemispheres of male mice.** Radial plots showing (**A**) immune cell populations, reported as percentages of $CD45^+$ cells, and (**B**) T cell subsets, reported as percentages of $CD3^+$ cells at days 7 and 21 post-implantation. *Gran*, granulocytes; *MDM*, monocyte-derived macrophages; *DC*, conventional dendritic cells; *Mono*, monocytes; *NK*, natural killer cells; *EM*, effector memory; *CM*, central memory.

**Fig. S4**

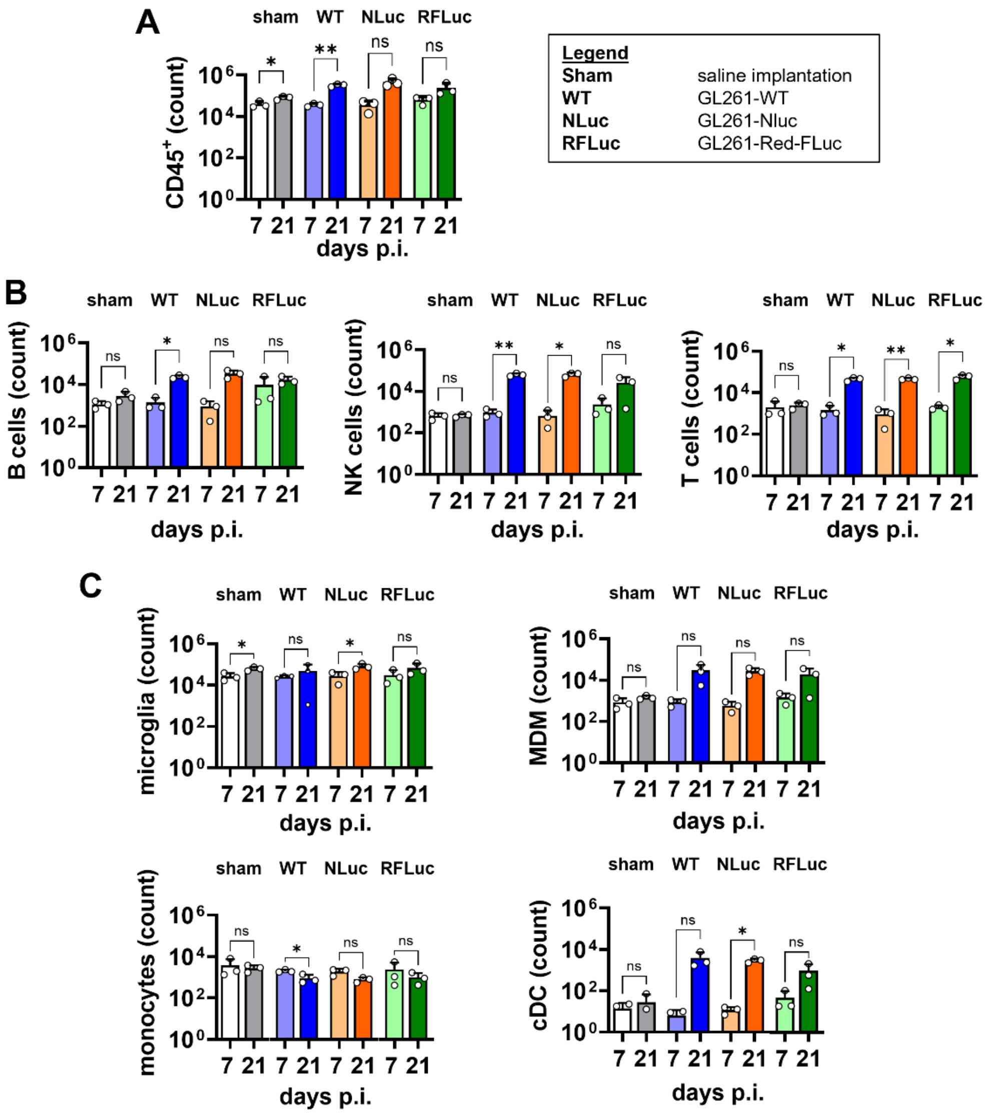


**Fig. S4. Immune cells in tumor-bearing brain hemispheres of male mice after GL261 implantation.** Absolute counts of (**A**) total CD45$^+$ immune cells, (**B**) lymphoid immune cells, and (**C**) myeloid immune cells brains of mice implanted with either GL261-WT (*WT*), GL261-NLuc (*NLuc*), or GL261-Red-FLuc (*RFLuc*) cells or saline (*sham*) at days 7 and 21 post-implantation (p.i.). Data are presented as means ± SD from $n = 3$ mice per group from one study cohort. Data were compared between days 7 and 21 by unpaired $t$ test. *$p < 0.05$; **$p < 0.005$; ***$p < 0.001$; *ns*, not significant. *NK*, natural killer cells; *MDM*, monocyte-derived macrophages; *cDC*, conventional dendritic cells.

**Fig. S5**

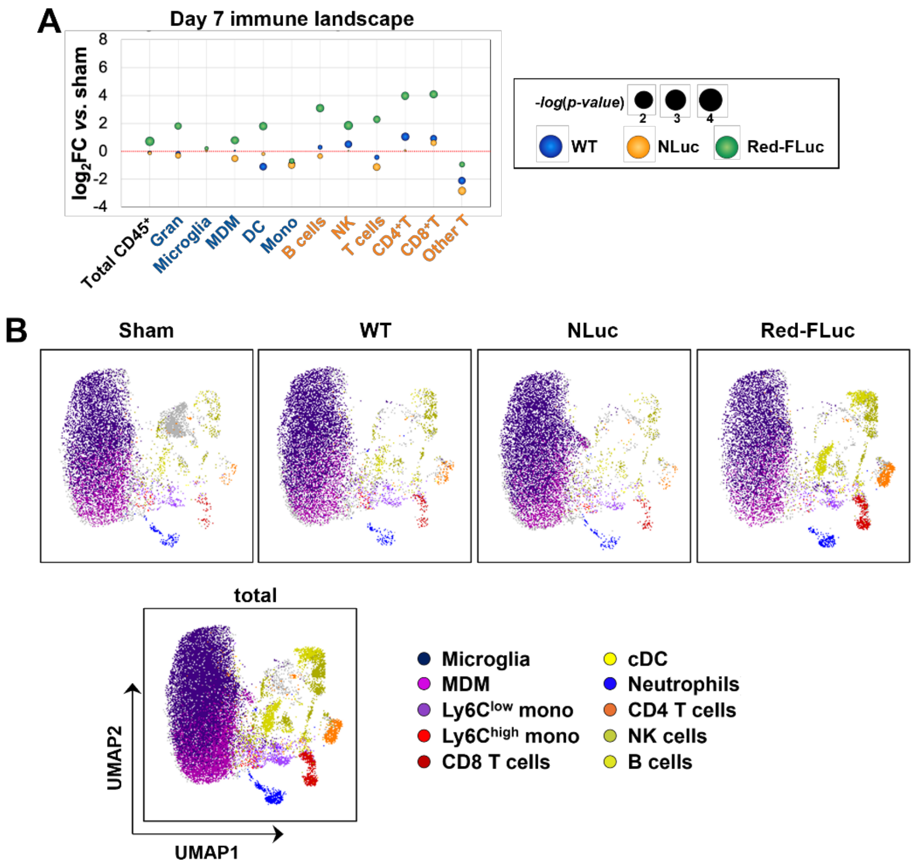


**Fig. S5. Immune cell landscape in tumor-bearing brain hemispheres of male mice.** (**A**) Bubble plot of immune cell composition at day 7 post-implantation (p.i.), shown as $\log_2$ fold change (FC) in cell counts relative to sham-implanted controls. Bubble size is inversely proportional to the corresponding *p* value. (**B**) UMAP visualization of immune cell profiles at day 7 p.i. where cell types were assigned using the manual gating strategy shown in Fig. S2. *Gran*, granulocytes; *MDM*, monocyte-derived macrophages; *cDC*, conventional dendritic cells; *Mono*, monocytes; *NK*, natural killer cells.

**Fig. S6**

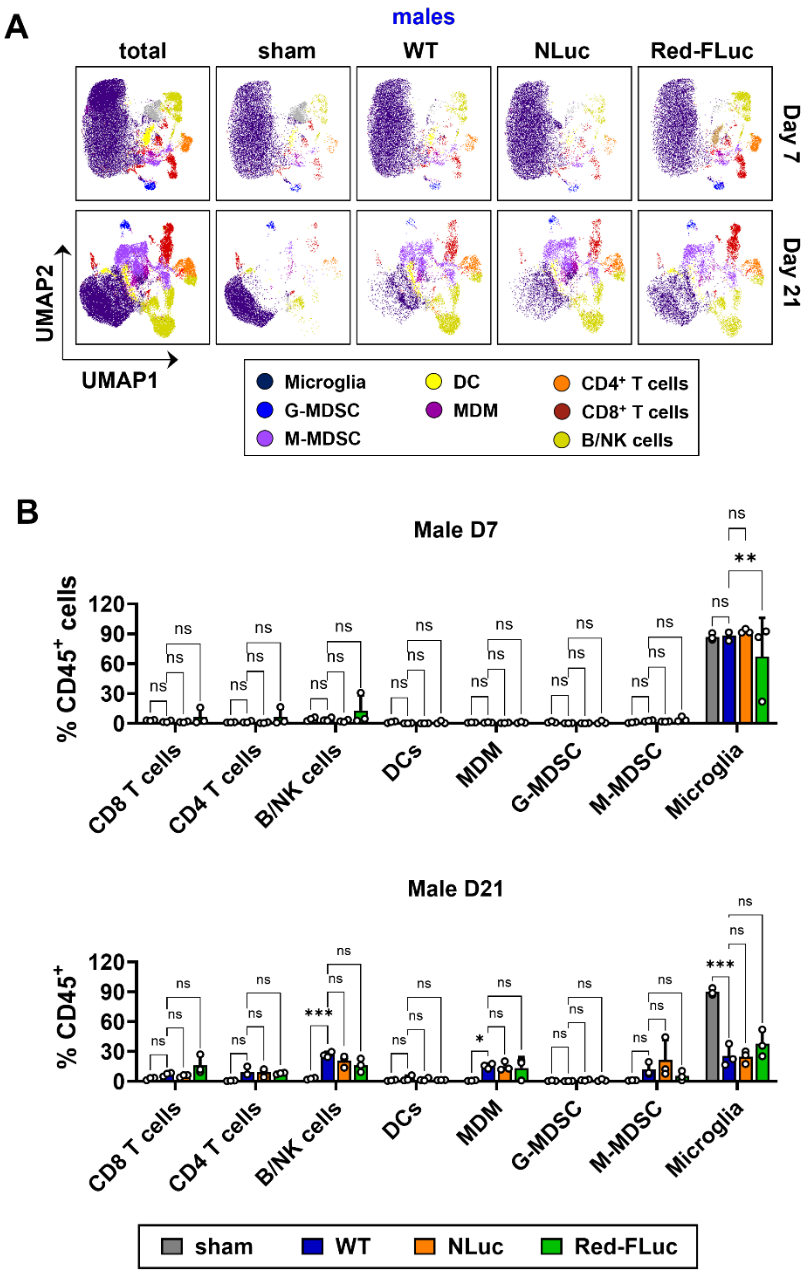


**Fig. S6. FlowSOM analysis of immune cell populations in tumor-bearing brain hemispheres of male mice.** (**A**) UMAP visualization of immune cell profiles at days 7 and 21 post-implantation using unsupervised FlowSOM clustering in FlowJo. (**B**) FlowSOM-based quantification of relative immune cell numbers across experimental groups. Data are shown as means ± SD from 3 mice per group from representative study cohort and compared by Welch's ANOVA. *$p < 0.05$; **$p < 0.005$; ***$p < 0.001$; ns, not significant. *G-MDSC*, granulocytic myeloid-derived suppressor cells; *M-MDSC*, monocytic myeloid-derived suppressor cells; *DC*, dendritic cells; *MDM*, monocyte-derived macrophages; *NK*, natural killer cells.

**Fig. S7**

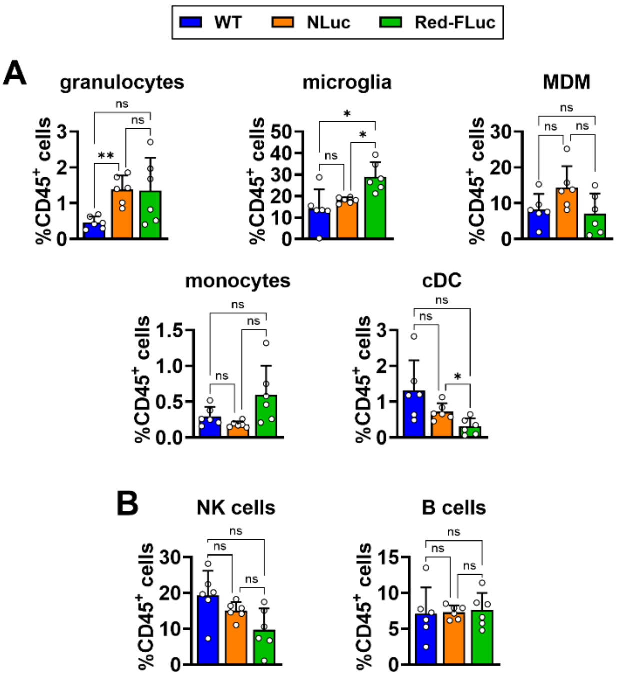


**Fig. S7. Immune cell counts in tumor-bearing brain hemispheres of male mice at day 21 post-implantation.** Relative abundance of (**A**) myeloid; and (**B**) lymphoid immune cells in brains of mice implanted with either GL261-WT (*WT*), GL261-NLuc (*NLuc*), or GL261-Red-FLuc (*Red-FLuc*). Data are presented as mean ± SD of 6 mice per group, and means were compared using Welch's ANOVA with Dunnett's correction for multiple comparisons. **p* < 0.05; ***p* < 0.005; ****p* < 0.001; ns, not significant. *MDM*, monocyte-derived macrophages; *cDC*, conventional dendritic cells; *NK*, natural killer cells.

**Fig. S8**

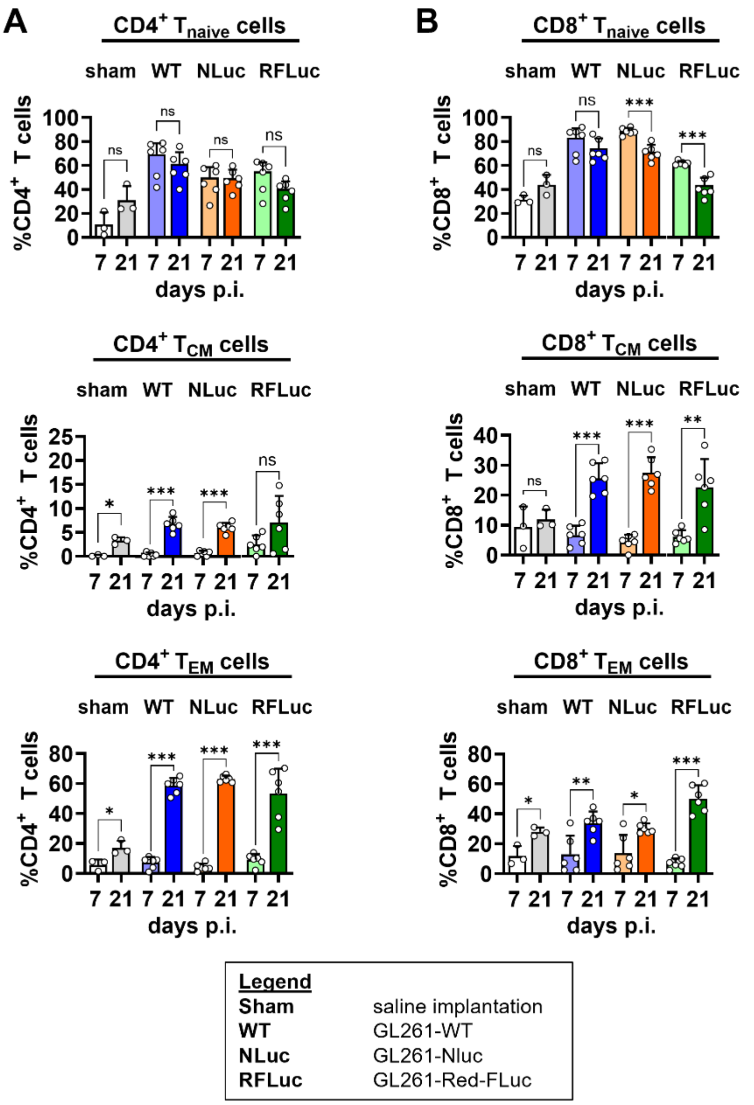


**Fig. S8. T cell subsets in tumor-bearing brain hemispheres of male mice.** Relative abundance of naïve T cells ($T_{naive}$), effector memory T cells ($T_{EM}$), and central memory T cells ($T_{CM}$) among (**A**) $CD4^+$ T cells; and (**B**) $CD8^+$ T cells in brains of male mice implanted with either saline control (*sham*), GL261-WT (*WT*), GL261-NLuc (*NLuc*), or GL261-Red-FLuc (*RFLuc*) cells. Data are presented as mean ± SD from 6 mice per group, and means were compared between day 7 and day 21 post-implantation (p.i.) by Welch's *t* test. *$p < 0.05$; **$p < 0.005$; ***$p < 0.001$; *ns*, not significant.

**Fig. S9**

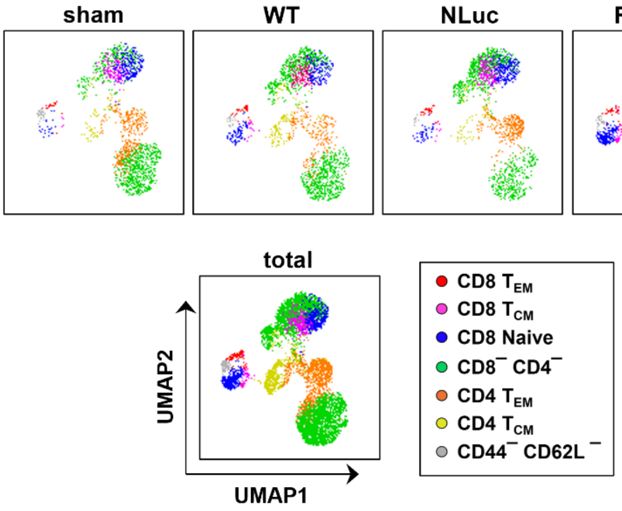


**Fig. S9. T cell landscape in tumor-bearing brain hemispheres of male mice at day 7 post-implantation.** UMAP visualization of T cell profiles in male mice implanted with either saline control (*sham*), GL261-WT (*WT*), GL261-NLuc (*NLuc*), or GL261-Red-FLuc (*Red-FLuc*) cells at day 7 post-implantation. UMAPs were generated from FCM data gated manually according to the strategy in Fig. S2.

**Fig. S10**

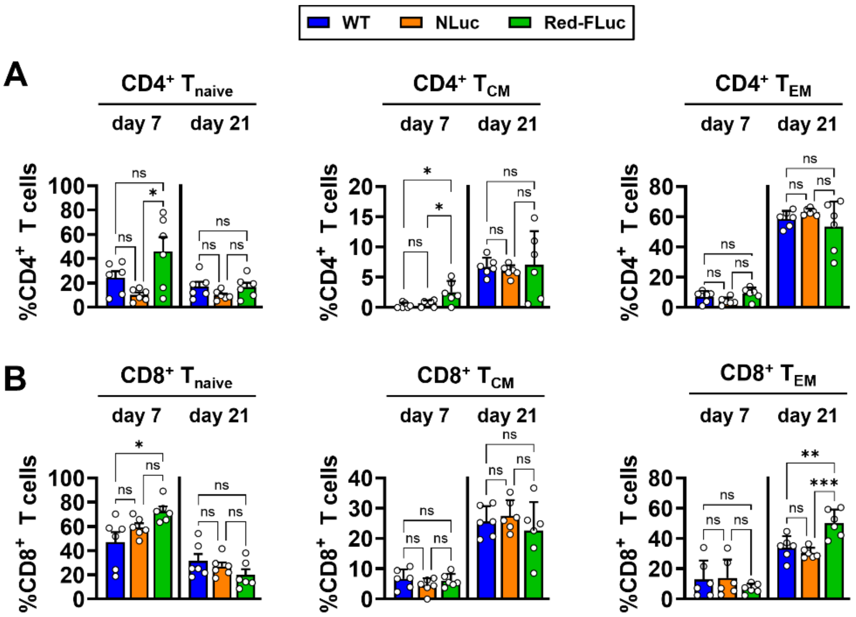


**Fig. S10. T cell subsets in tumor-bearing brain hemispheres of male mice.** Relative abundance of naïve T cells ($T_{naive}$), effector memory T cells ($T_{EM}$), and central memory T cells ($T_{CM}$) among (**A**) CD4$^+$ T cells; and (**B**) CD8$^+$ T cells in brains of male mice implanted with either GL261-WT (*WT*), GL261-NLuc (*NLuc*), or GL261-Red-FLuc (*Red-FLuc*) cells. Data are presented as mean ± SD from 6 mice per group, and means were compared by ordinary one-way ANOVA with Tukey's correction for multiple comparisons. *$p < 0.05$; **$p < 0.005$; ***$p < 0.001$; *ns*, not significant.

**Fig. S11**

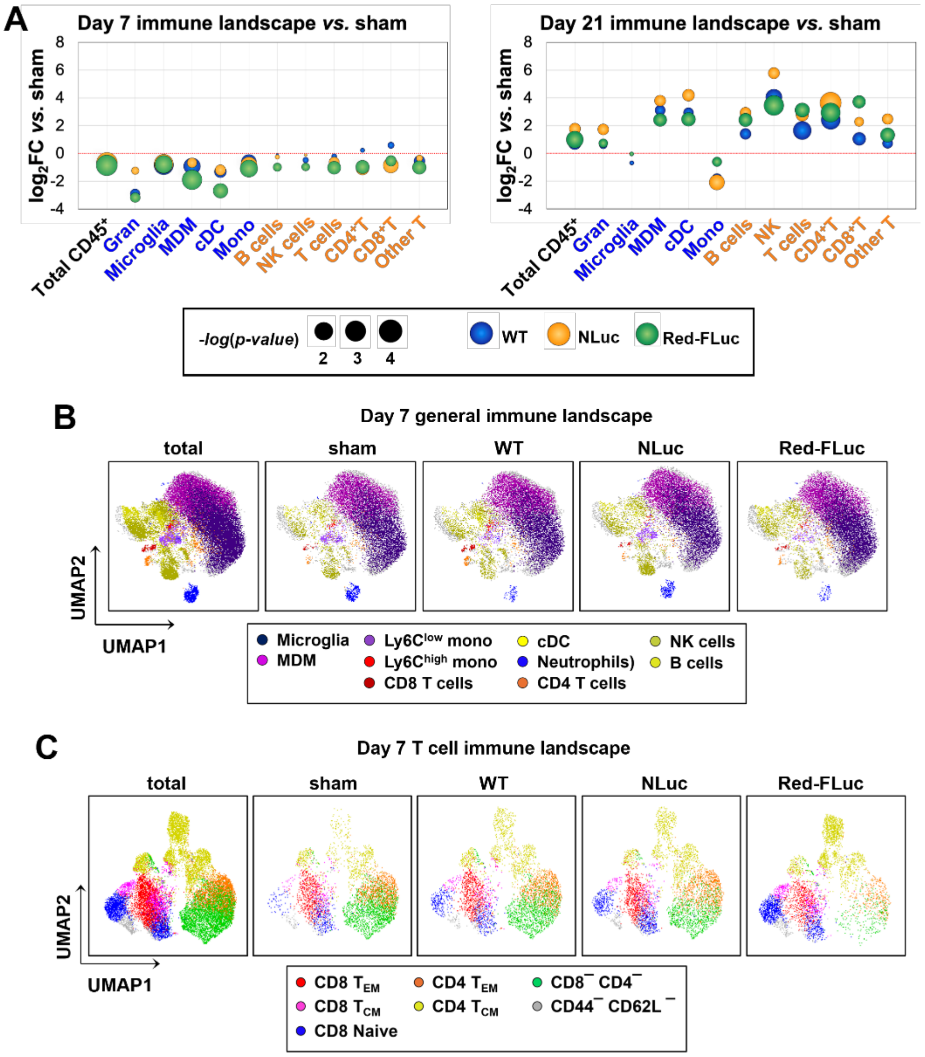


**Fig. S11. Immune and T cell landscapes in tumor-bearing brain hemispheres of female mice at day 7 post-implantation.** (**A**) Bubble plot of immune cell composition at day 7 post-implantation (p.i.), shown as $\log_2$ fold change (FC) in cell counts relative to sham-implanted controls. Bubble size is inversely proportional to the corresponding *p* value. (**B** and **C**) UMAP visualization of general immune (**B**) and T cell (**C**) profiles at day 7 p.i. UMAPs were generated from manually gated data using the strategy in Fig. S2. *Gran*, granulocytes; *MDM*, monocyte-derived macrophages; *cDC*, conventional dendritic cells; *Mono*, monocytes; *NK*, natural killer cells; $T_{CM}$, central memory T cells; $T_{EM}$, effector memory T cells.

**Fig. S12**

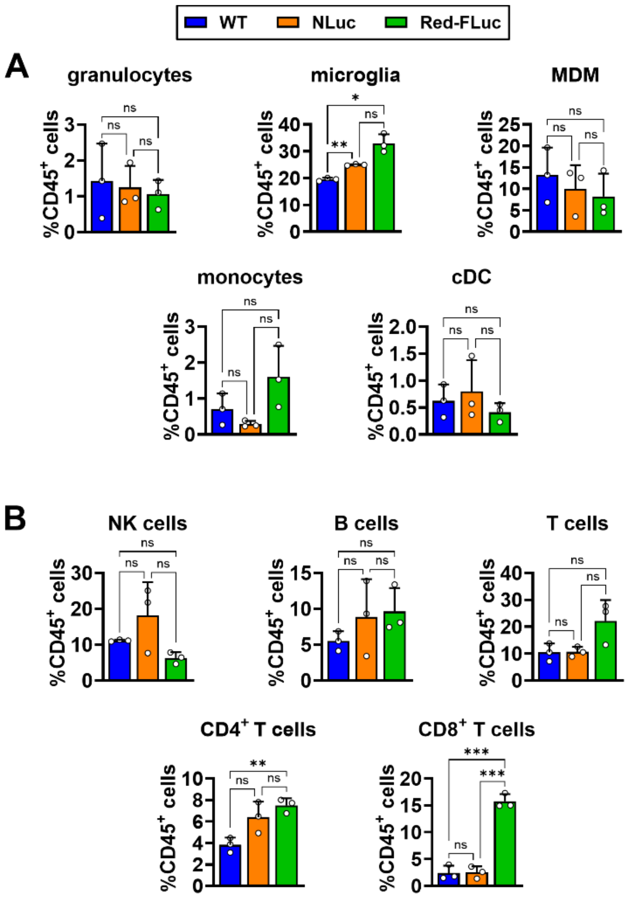


**Fig. S12. Relative abundance of immune cells in tumor-bearing brain hemispheres of female mice at day 21 post-implantation.** Relative counts of (**A**) myeloid cells and (**B**) lymphoid cells in tumor-bearing (TB) brain hemispheres from mice implanted with either GL261-WT (*WT*), GL261-NLuc (*NLuc*), or GL261-Red-FLuc (*Red-FLuc*) cells, shown as percentages of $CD45^+$ cells. Data are means ± SD from 3 mice per group and were analyzed by one-way ANOVA with Tukey's correction for multiple comparisons. *$p < 0.05$; **$p < 0.005$; ***$p < 0.001$; *ns*, not significant. *MDM*, monocyte-derived macrophages; *cDC*, conventional dendritic cells; *NK*, natural killer cells.

**Fig. S13**

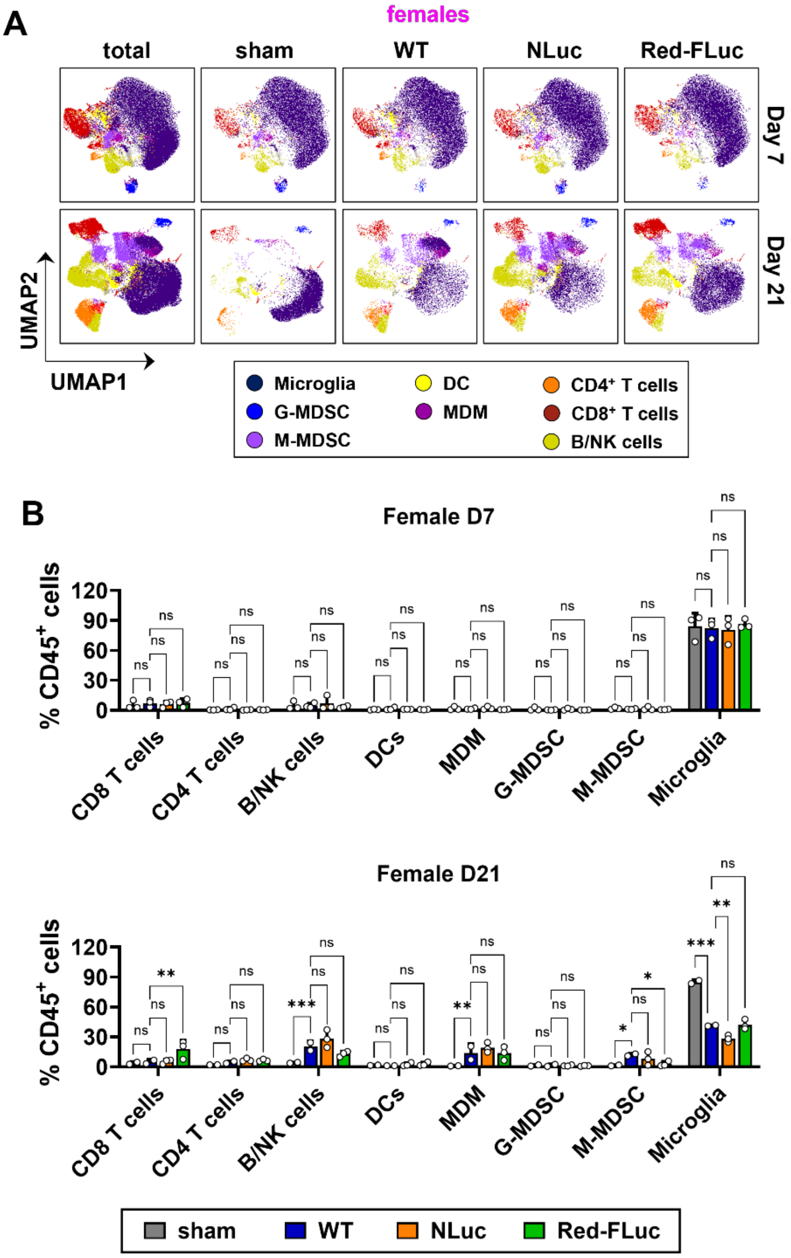


**Fig. S13. FlowSOM analysis of immune cell populations in tumor-bearing brain hemispheres of female mice.** (**A**) UMAP visualization of immune cell profiles in mice implanted with saline control (*sham*), GL261-WT (*WT*), GL261-NLuc (*NLuc*), or GL261-Red-FLuc (*Red-FLuc*) cells at days 7 and 21 post-implantation using unsupervised FlowSOM clustering. (**B**) FlowSOM-based quantification of relative immune cell numbers across experimental groups. Data are shown as mean ± SD from 3 mice per group and means were compared by two-way ANOVA with Holm-Šídák correction for multiple comparisons. *$p < 0.05$; **$p < 0.005$; ***$p < 0.001$; ns, not significant. *G-MDSC*, granulocytic myeloid-derived suppressor cells; *M-MDSC*, monocytic myeloid-derived suppressor cells; *DC*, dendritic cells; *MDM*, monocyte-derived macrophages; *NK*, natural killer cells.

**Fig. S14**

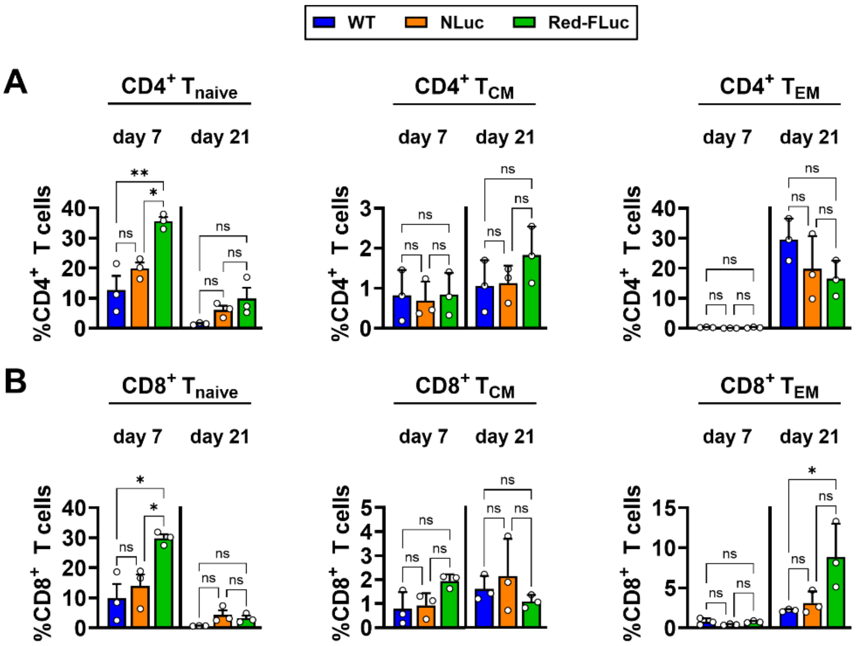


**Fig. S14. T cell subset distribution in tumor-bearing brain hemispheres of female mice.** Relative abundance of naïve T cells ($T_{naive}$), effector memory T cells ($T_{EM}$), and central memory T cells ($T_{CM}$) among (**A**) total $CD4^+$ T cells; and (**B**) total $CD8^+$ T cells from female brains implanted with either GL261-WT (*WT*), GL261-NLuc (*NLuc*), or GL261-Red-FLuc (*Red-FLuc*) cells. Data are presented as mean ± SD from 3 mice per group and means were compared by ordinary one-way ANOVA with Tukey's correction for multiple comparisons. *$p < 0.05$; **$p < 0.005$; ***$p < 0.001$; ns, not significant.

**Fig. S15**

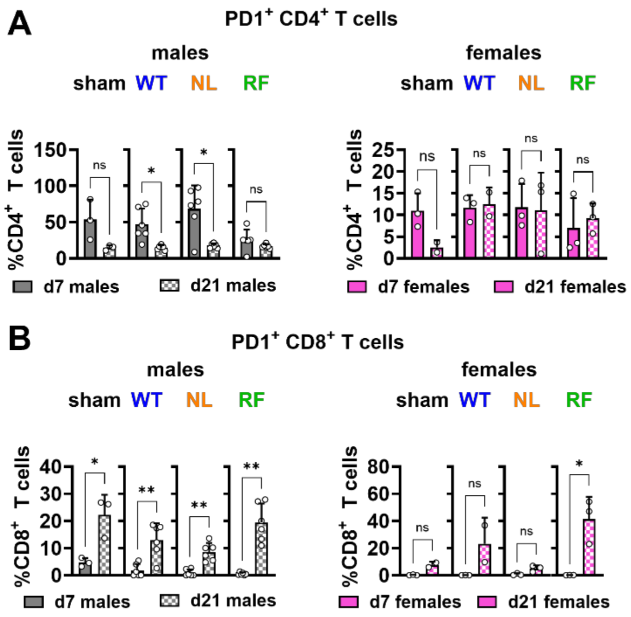


**Fig. S15. Temporal dynamics of PD1 expression prevalence on T cells in tumor-bearing brain hemispheres.** Frequencies of (**A**) $PD1^+$ $CD4^+$ T cells and (**B**) $PD1^+$ $CD8^+$ T cells in male and female mice implanted with either saline control (*sham*), GL261-WT (*WT*), GL261-NLuc (*NL*), or GL261-Red-FLuc (*RF*) cells at days 7 and 21 post-implantation. Data are shown as mean ± SD of 6 mice per group in male mice and 3 mice per group in female mice. Means were compared between day 7 and day 21 by Welch's *t* test. *$p < 0.05$; **$p < 0.005$; ***$p < 0.001$; *ns*, not significant.

**Fig. S16**

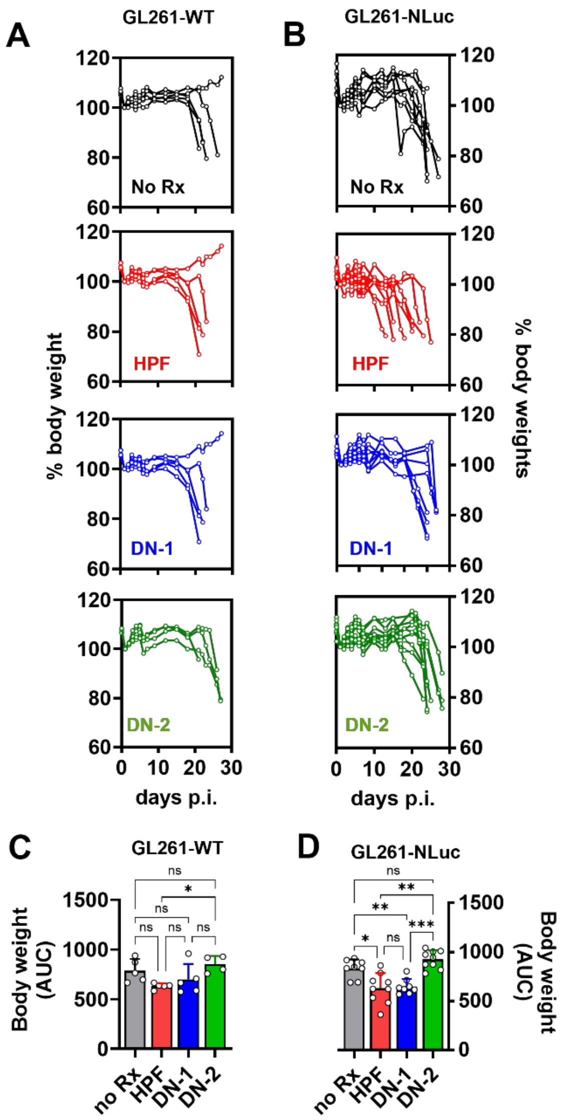


**Fig. S16. Body weight changes in mice implanted with tumors pre-treated with Zika virus live-attenuated vaccines (ZIKV-LAV).** (**A** and **B**) Body weight changes in mice implanted intracranially with GL261-WT (**A**) or GL261-NLuc (**B**) cells that were either mock-treated or pre-infected with ZIKV HPF or ZIKV-LAV (DN-1 and DN-2) strains 24h before implantation. Body weight is expressed as a percentage of weight measured at day 1 post-implantation. (**C** and **D**) Area under the curve (AUC) analysis of the body weight curves shown in (**A**) and (**B**), respectively. Data are presented as mean ± SD and were compared using Welch's ANOVA with Dunnett's correction for multiple comparisons. *$p < 0.05$; **$p < 0.005$; ***$p < 0.001$; ns, not significant.

**Fig. S17**

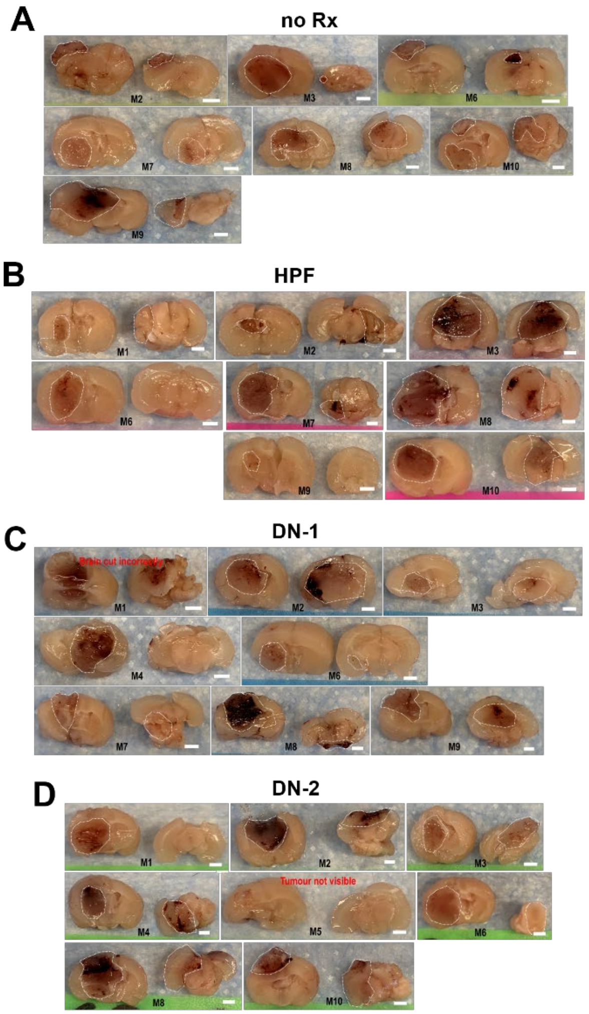


**Fig. S17. Gross pathology of tumor-bearing brains after implantation with ZIKV-treated GL261-NLuc cells.** Representative fixed coronal brain sections from mice implanted intracranially with GL261-NLuc cells that were mock-treated (No Rx; **A**) or pre-infected for 24 hours with HPF (**B**), DN-1 (**C**), or DN-2 (**D**) before implantation. Brains were harvested when mice reached the humane endpoint. Tumor margins are indicated by white dashed lines. Scale bar = 2 mm.
